# Feedback control via H3K9 methylation and siRNA biogenesis autonomously maintains RNAi heterochromatin through continuous low-level transcription

**DOI:** 10.64898/2026.10.28.762611

**Authors:** Sumanth Kumar Maheshwaram, Amith Zafal Abdulla, Serena Chiu, Dionysus Morris-Evans, Jane Kondev, Kaushik Ragunathan

## Abstract

The logic of RNA-based silencing involves nascent RNA molecules that serve as scaffolds to recruit proteins that repress transcription. Despite transcription and silencing being antagonistic, RNA-based recruitment mechanisms have been widely adopted in organisms ranging from yeast to humans. In fission yeast (*Schizosaccharomyces pombe*), the nuclear RNAi pathway that establishes heterochromatin at pericentromeric repeats to silence transcription exemplifies this paradox. Current models propose that cells reconcile these opposing activities by confining pericentromeric transcription to S phase while maintaining silencing during the rest of the cell cycle. Here, using strand-specific single-molecule RNA FISH (smFISH), we detect forward and reverse pericentromeric transcripts with high sensitivity and single-cell resolution. We find that repeat-associated transcription is rare, heterogeneous and happens independently of cell-cycle control. Instead, transcript production is governed solely by feedback involving small RNA biogenesis and H3K9 methylation. By combining mathematical modeling with experiments, we show that low levels of H3K9me or siRNAs prevent heterochromatin establishment, whereas high levels of H3K9me cause stochastic switching. Our work reveals that the same RNA-based feedback architecture can sample distinct transcriptional regimes, with feedback strength determining how often cells produce transcripts without any external regulatory inputs.

## Introduction

The regulation of gene expression in eukaryotic cells is governed at least in part by the covalent modification of DNA and DNA packaging proteins called histones ^1^. Histone and DNA modifications mark distinct genomic loci to establish specialized chromatin states that can activate or silence gene expression ^2^. H3K9 methylation is a repressive modification that is enriched at transcriptionally silent regions called heterochromatin, which includes pericentromeric repeats, telomeres and transposable elements in organisms such as yeast, plants and humans ^3,4^. Another histone modification, H3K27 methylation, marks developmental genes such as *Hox* gene clusters in flies and mammals ^5–7^. Since enzymes that silence gene expression have no intrinsic affinity for particular DNA sequences, sequence-specific recognition factors must guide histone modifying enzymes to distinct genomic loci ^8^. Sequence-specific recruitment factors fall into two major classes: the first involves proteins that recognize and bind DNA ^9–11^ and the second consists of proteins that bind to nascent RNA transcribed from the target locus ^12^.

While it is relatively straightforward to envision how DNA binding proteins recognize specific sequence motifs and recruit effectors, the logic of RNA based silencing presents a paradox. Transcription must precede silencing to produce nascent RNA transcripts that serve as scaffolds to recruit silencing factors that repress transcription (**Figure 1A**). However, transcription leads to nucleosome disassembly, histone acetylation, and the loss of chromatin compaction, which antagonize heterochromatin establishment ^13,14^. Despite these two processes being at odds with each other, organisms such as yeasts and protozoans (small RNA mediated silencing) ^15,16^, plants (RNA dependent DNA methylation) ^17–19^, flies (piRNA mediated silencing) ^20,21^, and mammals (PRC2 and HUSH mediated silencing) ^22–24^ have repeatedly adopted RNA based silencing strategies highlighting the versatility of this targeting system. However, we do not understand how these systems have evolved to ensure that transcription can co-exist in tandem with silencing.

**Figure 1.**
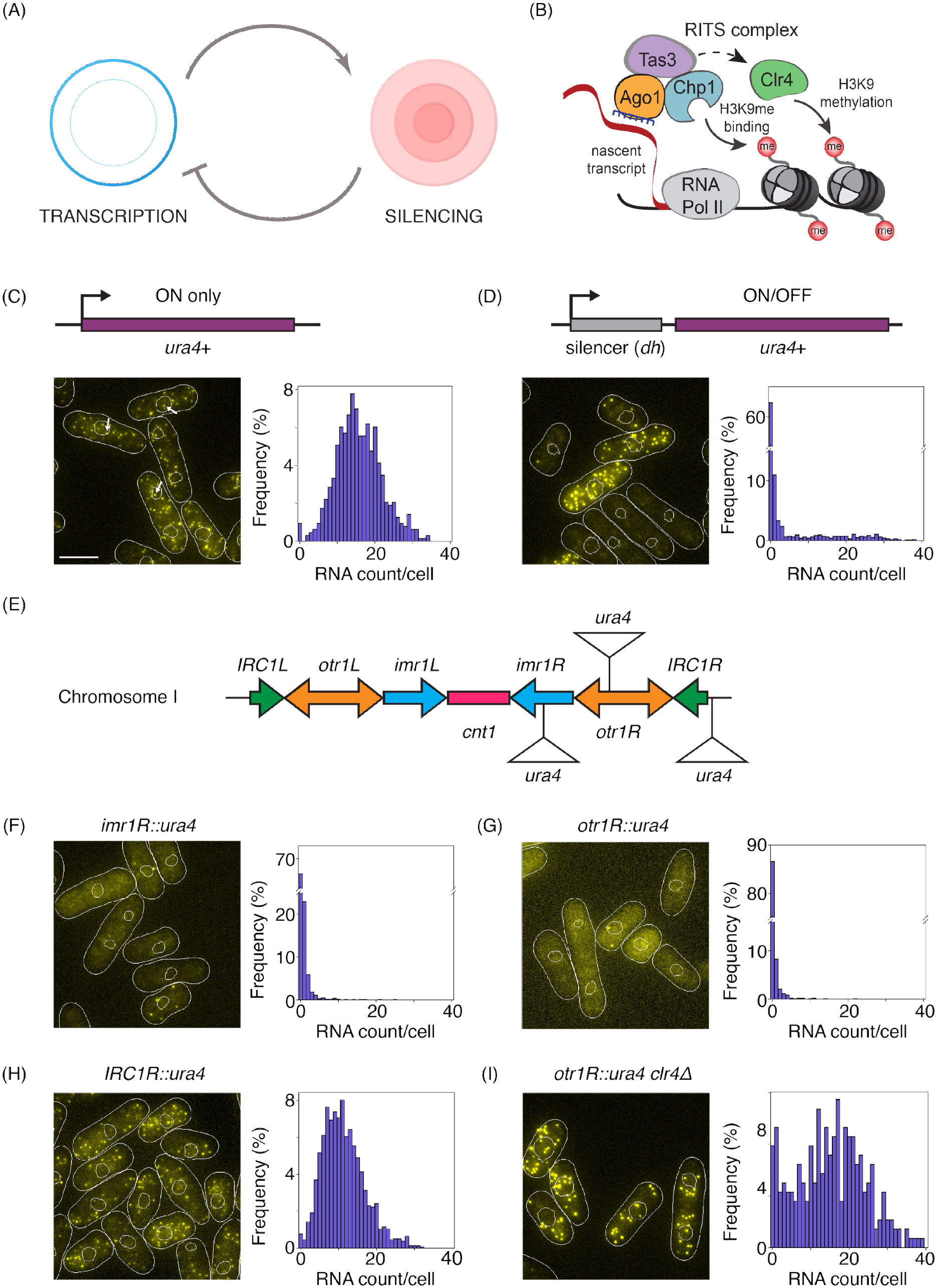
Single molecule RNA FISH (smFISH) detects individual *ura4+* coding transcripts that originate from endogenous loci or from pericentromeric repeats. **(A)** RNA-based silencing systems present a paradox where the silencing of a region requires its own transcription. **(B)** Heterochromatin maintenance at pericentromeres in *S*.*pombe* is achieved by RNAi which involves bivalent recognition of H3K9me3 on chromatin and the nascent transcripts generated from the same region. The specific recruitment of RITS complex occurs through the help of siRNA bound to RITS complex and hybridizes to RNA. This results in a cascade of downstream interactions leading to establishment and stable maintenance of silencing. **(C)** Fluorescence microscopy images of endogenous *ura4+* transcripts using single-molecule RNA-FISH. White arrows are pointing towards the nascent transcripts. Histogram of RNA counts per cell are shown to the right. Scale bar is 5 μm in all the images. **(D)** Fluorescence microscopy images of endogenous *ura4+* transcripts when *dh* sequence is inserted in the *ura4+* locus. **(E)** Schematic showing pericentromeric repeats in the chromosome I. **(F)** Fluorescence microscopy images of *ura4+* transcripts when *ura4+* gene is inserted at the pericentromeric *imr1R* locus (*imr1*R::*ura4+*), with histogram of RNA counts/cell to the right. (**G-I**) Similar to (F) for *otr1*R::*ura4+* (G), *irc1*R::*ura4+* (H), *otr1R*::*ura4+ clr4Δ* (I).

The fission yeast *Schizosaccharomyces pombe* represents an outstanding model system to investigate the logic of RNA based silencing mechanisms ^16^ (**Figure 1B**). At pericentromeric *dg* and *dh* repeats, RNA polymerase II produces non-coding transcripts that are converted into double-stranded RNA by the RNA-directed RNA polymerase complex (RDRC) and processed into siRNAs by Dcr1 ^25–27^. These siR-NAs are bound by Ago1 which is part of a three-protein complex called RNA-induced initiation of transcriptional silencing (RITS) complex ^28^. The RITS complex is tethered to nascent transcripts through complementary small RNA dependent base-pairing interactions and to chromatin through recognition of pre-existing H3K9me ^29^. The RITS complex recruits the H3K9 methyltransferase, Clr4 to pericentromeric repeats where it catalyzes and spreads H3K9me ^30,31^. Ultimately, H3K9me promotes transcriptional gene silencing which leads to the exclusion of RNA polymerase II from pericentromeric repeats. Deleting either the small RNA binding protein Argonaute (*ago1Δ)* or the H3K9 methyltransferase Clr4 (*clr4Δ)* leads to the loss of transcriptional silencing, H3K9me and siRNA biogenesis ^25,27,32^. Hence, in fission yeast nuclear RNAi exemplifies the core logic of RNA based silencing systems where transcription is required to maintain stable silencing.

The current model which explains how cells navigate the conflict between transcription and silencing proposes that the two processes are temporally insulated from each other ^33,34^. According to this model, pericentromeric repeats are largely silent through most of the fission yeast cycle (G2/M/G1) but are transcribed within a narrow window co-incident with S phase. During this period, H3S10 phosphorylation leads to HP1 dissociation (Swi6 in *S*.*pombe*) thus creating an environment that is permissive for transcription ^35,36^. S-phase restricted nascent transcripts subsequently serve as scaffolds to nucleate and re-establish heterochromatin once every cell cycle ^37,38^. Nevertheless, evidence in support of this model is based on cell synchronization approaches that either disrupt cellular physiology ^34,39^ or compromise heterochromatin stability ^33^. Moreover, population-level transcription measurements average across millions of cells, which can obscure transcriptional events that are rare or heterogeneous with distinct temporal dynamics across individual cells ^40–42^.

In this study we address this conceptual gap with respect to how and when pericentromeric repeats are transcribed using a strand-specific single-molecule fluorescence *in situ* hybridization (smFISH) approach ^43–45^. This enables us to directly count nascent RNA transcripts with high sensitivity and resolution in individual cells without the need for any cell synchronization methods. We show that both forward and reverse non-coding RNA strands associated with pericentromeric repeat sequences are transcribed across all phases of the cell-cycle as opposed to being restricted to the S-phase of the cell-cycle. Based on our results, we propose that continuous, low-levels of transcription are required for stable heterochromatin with transcript levels being autonomously regulated through feedback involving H3K9 methylation and siRNA biogenesis.

## Results

### Single molecule RNA FISH (smFISH) detects individual *ura4+* coding transcripts that originate from endogenous loci or from pericentromeric repeats

We implemented a single-molecule FISH (smFISH) approach to detect mature and nascent transcripts in *S*.*pombe* cells ^43–45^. smFISH uses several fluorescent oligonucleotide probes that simultaneously hybridize to individual transcripts, labeling both nascent transcripts at the site of transcription and mature transcripts within the cytoplasm. Since transcripts can be detected with single molecule and single cell resolution, we can acquire information about transcription dynamics without the need for any cell synchronization approaches.

To establish a baseline for the sensitivity and resolution of smFISH, we first labeled *ura4+* transcripts produced from its endogenous locus, where it is constitutively transcribed, and compared its localization and abundance to those of *ura4+* transcribed when a silencer is inserted next to it. The silencer represents a 2.8 kb sequence derived from the pericentromeric repeats (*dh*) that induces RNAi-mediated heterochromatin establishment ^46^. Our measurements show that endogenous *ura4+* transcription resulted in about ∼15 RNA molecules per cell (**Figure 1C**), with only 1% of cells having no transcripts (see **Supplementary Figures 1A-E** for details about the analysis workflow). This observation is consistent with transcription being stochastic ^47–49^.

In contrast, placing a silencer adjacent to *ura4+* produced two distinct populations. About 80% of cells had few to no transcripts (≤ 3), consistent with *ura4+* being silenced, while the remaining cells exhibited high levels of transcription (∼ 20 transcripts) with large cell-to-cell variance in transcript abundance (**Figure 1D**). While the majority of *ura4+* transcripts in both instances were localized within the cytoplasm, we frequently detected RNA molecules within the nucleus that correspond to sites of nascent transcription (white arrow, **Figure 1C**).

Next, we used classical heterochromatin reporter strains in which *ura4+* is inserted at distinct locations within centromere 1 (*cen1*) (**Figure 1E**). Previous studies have shown that cells with *ura4+* insertions within the outer (*otr1R::ura4+*, **Figure 1G**) or innermost (*imr1R::ura4+*, **Figure 1F**) ^50,51^ repeats are resistant to 5-fluoroorotic acid (5-FOA) and fail to grow on media lacking uracil (-URA) due to silencing. Indeed, we detected no transcripts in the majority of cells (∼85% of cells in *otr1R::ura4+* and ∼65% of cells in *imr1R::ura4+*). The small fraction of cells in which we detected transcripts, averaged only ∼1-3 RNA molecules per cell across both *otr1R::ura4+* or *imr1R::ura4+* insertions. However, inserting the same *ura4+* sequence outside pericentromeric heterochromatin boundaries (*IRC1R::ura4+*, **Figure 1H**) ^52^ led to a significant increase in transcript numbers (∼11 transcripts/cell). These results confirm that our smFISH measurements accurately capture position effects associated with *ura4+* expression within pericentromeric repeats.

To confirm that *ura4+* is epigenetically silenced, we deleted the H3K9 methyltransferase Clr4 (*clr4Δ*, **Figure 1I**). The subsequent loss of silencing led to a dramatic increase in *ura4+* transcription with a mean value of ∼15 transcripts per cell. We observed similar results upon deleting the small RNA binding protein, Ago1 (*ago1Δ*) which captures the strong dependence of pericentromeric silencing on RNAi (**Supplementary Figure 1F**). In summary, we confirmed that smFISH detects transcripts with single-molecule sensitivity, enabling us to count from as few as one to several tens of transcripts in individual cells.

### Pericentromeric non-coding transcripts are rare and preferentially retained within the nucleus

There are approximately 20 repeats of *dh* and 14 repeats of *dg* across three chromosomes which serve as scaffolds to nucleate heterochromatin assembly ^53–56^ (See **Supplementary Note 1** for detailed analysis of repeat sequence similarity). Unlike coding transcripts, non-coding pericentromeric repeats are transcribed in a bidirectional manner ^25,26,57^. To simultaneously measure bidirectional transcription and how it contributes to heterochromatin establishment, we imaged forward and reverse *dg* and *dh* transcripts using two distinct fluorophores. Our smFISH approach is mostly insensitive to nucleotide-level differences between repeats, and therefore our measurements capture the net transcriptional output across similar *dg* or *dh* sequences.

Upon labeling either *dg* or *dh* repeats, we observed that wild-type cells produce either no transcripts or as few as 1-3 transcripts per cell in the forward (*dgF* and *dhF)* and reverse (*dgR* and *dhR)* direction (**Figure 2A,B and Supplementary Figure 2A,B**). We confirmed that our measurements detect RNA and not DNA since pre-treating cells with RNAse leads to the complete elimination of all measurable hybridization-dependent signal (**Supplementary Figure 2C,D**). We also expected that non-coding RNA transcripts that originate from pericentromeric repeats will co-localize with the histone H3 variant, CENP-A that marks centromeres ^58^. We imaged Cnp1-GFP in conjunction with *dg* forward (*dgF*) and reverse (*dgR*) transcripts and observed colocalization in 25% and 27% of cells, respectively (**Figure 2G**). These percentages include cells without detectable nascent transcripts (**Figure 2B**). Similarly, Cnp1-GFP colocalizes with 25% of *dh* forward (*dhF*) and 41% of *dh* reverse (*dhR*) transcripts (**Figure 2G**). The co-localization with CENP-A suggest that the signals we measure correspond to *bona fide* pericentromeric transcripts rather than spurious localization errors.

**Figure 2.**
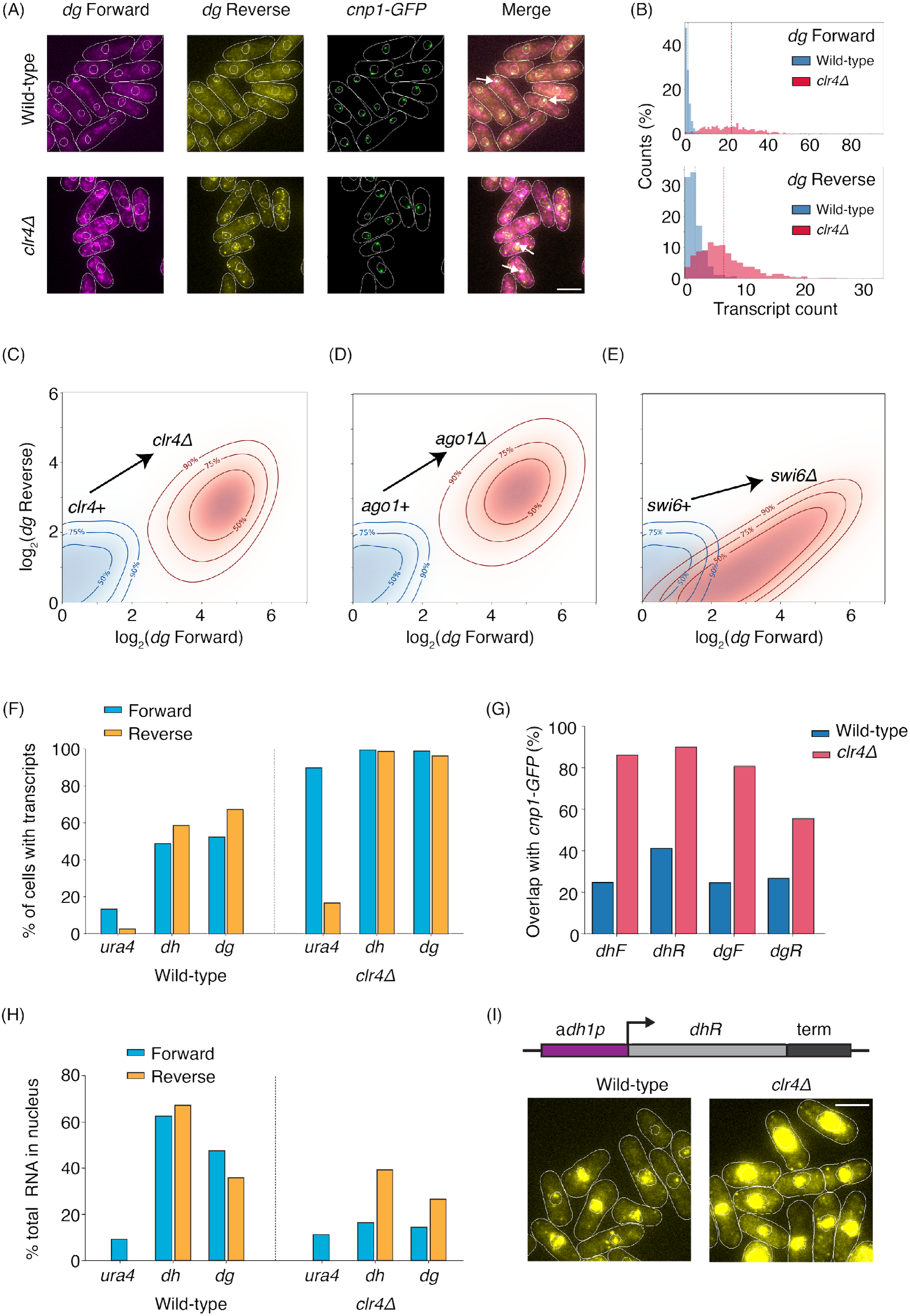
Pericentromeric non-coding transcripts are rare and preferentially retained within the nucleus. **(A)** Top panel shows fluorescence images of forward (magenta) and reverse (yellow) transcripts from *cen1 dg* repeat. Colocalization of transcripts with Cnp1-GFP confirms they are generated from the centromere. Bottom panel shows the fluorescence images of *dg* in *clr4Δ* mutant. Scale bar is 5 μm in all the images. **(B)** Histograms of *dg* forward and *dg* reverse RNA counts. **(C)** The contour plot shows increase in pericentromeric transcription in both directions in *clr4Δ* cells compared with wild-type. Blue and red contours represent the wild-type and the mutant cells respectively. **(D,E)** Similar contour for *ago1Δ* (D) and *swi6Δ* (E). **(F)** Percentage of cells with transcripts is quantified for forward and reverse transcripts of *ura4+, dg* and *dh*. **(G)** Colocalization statistics of Cnp1-GFP with *dg* and *dh* transcripts **(H)** Nuclear localization of forward (*dhF, dgF)* and reverse (*dhR, dgR*) RNA is quantified from single-cell images and compared with *ura4+*. **(I)** Fluorescence images of *dhR* transcripts when the *dhR* fragment driven by adh1 promoter is inserted at an ectopic locus.

Since pericentromeric *dg* and *dh* repeats are also silenced by H3K9me, deleting Clr4 (*clr4Δ*) led to a substantial increase in transcription across all repeats. Cells produce on average ∼22 transcripts per cell corresponding *to dgF* and ∼6 transcripts per cell corresponding to *dgR* (**Figure 2A-C,F**). We observed similar transcript numbers for *dh* repeats (*dhF*: 23, *dhR*: 7 transcripts/cell) (**Supplementary Figure 2G**). The colocalization of non-coding RNA sequences (in the forward and reverse direction) and CENP-A increased substantially in *clr4Δ* cells rising to 81% for *dgF* and 56% for *dgR* transcripts (87% with *dhF* and 90% with *dhR*) (**Figure 2G**). We observed a similar pattern of increased bidirectional transcription across all RNAi mutants including deletions of sub-units associated with the RITS complex (*ago1Δ*), RDRC (*rdp1Δ*) and Dicer (*dcr1Δ*) (**Figure 2D and Supplementary Figure 2F-H**). The only notable exception was deleting the HP1 protein, Swi6 (*swi6Δ*) (**Figure 2E, Supplementary Figure 2I**). In *swi6Δ*, we detected an asymmetric increase in transcription where forward-strand transcripts rise to levels comparable to *clr4Δ*, whereas reverse-strand transcripts remained close to what we typically measure in wild-type cells ^25^.. This leads to a skew in the ratio of *dgF* to *dgR* which is 7-fold in *swi6Δ* compared to being 4-fold in *clr4*Δ. These smFISH measurements including the observed asymmetry in *swi6*Δ match previous reports indicating that our labeling and detection methods recapitulate population-level trends associated with pericentromeric transcription.

### Nuclear retention of pericentromeric transcripts is strand- and repeat-specific

Our ability to directly image individual transcripts also provided insight into their spatial distribution (**Figure 2H**) ^45^. In wild-type cells, non-coding transcripts are mostly nuclear localized, implying that they are likely nascent transcripts. About 60% of *dhF* and 70% of *dhR* sequences are localized within the nucleus, in contrast to only 10% for *ura4+*. Hence, most protein coding *ura4+* transcripts, despite being inserted within pericentromeric repeats, are exported to the cytoplasm. To test if H3K9 methylation and heterochromatin establishment control these localization patterns, we quantified *dg* and *dh* transcript localization in *clr4Δ* cells. In *clr4Δ* cells, non-coding RNA transcripts corresponding to both forward sequences – *dgF* and *dhF* are exported to the cytoplasm and exhibit reduced nuclear retention when compared to both reverse sequences – *dgR* and *dhR*. This observation is even more striking when an ectopic *dhR* sequence is expressed using an *adh1* promoter in wild-type and *clr4Δ* cells where we observe substantial accumulation of this sequence within the nucleus (**Figure 2I**) ^46^. Hence, transcript localization patterns are both repeat and orientation specific (forward versus reverse) suggesting that some classes of non-coding RNAs (such as *dgR* and *dhR*) may have features that allow for their selective retention within the nucleus to initiate heterochromatin establishment.

### Pericentromeric transcription is cell-cycle independent and exhibits homeostatic scaling

Given our ability to measure single transcripts with high spatial resolution, we next tested when transcripts are produced during the *S*.*pombe* cell-cycle. To identify cells in S-phase, we expressed EGFP-Pcn1 (the PCNA ortholog in *S*.*pombe*) which exhibits a striking change in localization when cells replicate their DNA ^59,60^ (**Figure 3A,B**). Cells that have a diffuse PCNA distribution map to the G1, M or G2 phase whereas cells that have distinct PCNA foci are assigned to S-phase. We also measured cell length as an independent indicator of cell-cycle progression, since *S*.*pombe* cells continuously elongate until cytokinesis resets their length ^61–63^. We plotted a histogram of cell length versus the fraction of PCNA foci containing cells (**Figure 3C**). PCNA-foci containing cells (foci+) which are in S-phase have a bimodal distribution with cells enriched across both short and long cell lengths while PCNA-diffuse (foci-) cells which are in G2, M or G1 phases are enriched at intermediate lengths. This analysis recapitulates the expected relationship between cell length and cell-cycle stage, supporting our use of PCNA foci to identify cells in S phase.

**Figure 3.**
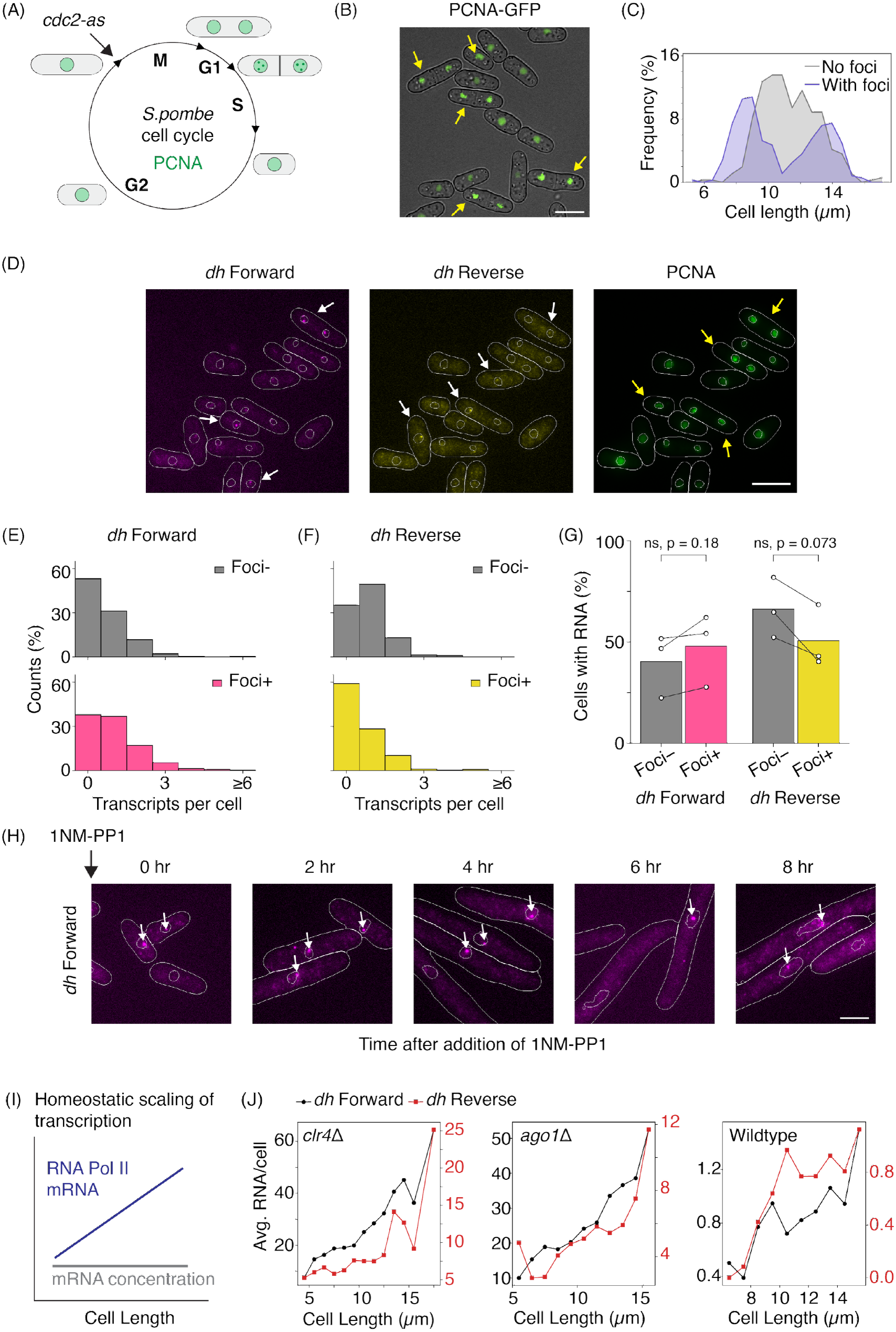
Pericentromeric transcription is cell-cycle independent and exhibits homeostatic scaling. **(A)** Schematic of the *S*.*pombe* cell cycle with G1, S, G2 and M phases. **(B)** Fluorescence image of live cells with (foci+) and without (foci-) PCNA-GFP foci. Yellow arrows point towards the foci+ cells and correlate with the cells having septum indicating they are in S-phase. Scale bar is 5 μm in all the images. **(C)** Cell length histograms for foci+ and foci-cells. Foci+ cells are enriched at the shorter and longer cell lengths whereas foci-cells are enriched at intermediate lengths. **(D)** Fluorescence images of *dh* forward and reverse transcripts in conjunction with PCNA-GFP. The white and yellow arrows point towards the foci- and foci+ cells respectively. **(E)** Histograms of *dh* forward transcript numbers per cell in foci- and foci+ cells indicate no cell cycle dependence. **(F)** Similar to (E) for *dh* reverse transcripts. **(G)** Percentage of cells with detectable *dh* RNA in foci- and foci+ cells . The data are from three independent replicates. **(H)** Fluorescence images of *dh* forward transcripts at 0hr, 2hr, 4hr, 6hr, 8hr timepoints after adding 1NM-PP1 to *cdc2-asM17* cells to arrest them at G2/M transition. **(I)** Schematic of transcript number scaling with cell size. **(J)** *dh* forward and reverse transcript numbers scale with cell size in *clr4Δ, ago1Δ* and wild-type cells.

Next, we imaged *dg* and *dh* transcripts in wild-type cells in conjunction with GFP-PCNA (**Figure 3D**). We plotted the distribution of transcripts based on PCNA foci status (foci+ versus foci–). PCNA foci status had no correlation with transcript numbers, i.e. cells which had foci (S-phase) had 1-3 transcripts which was similar to the distribution of transcripts that we observed in cell which had no foci (G2/M/G1 phase) (**Figure 3E-G, Supplementary Figures 3A-C**). Likewise, we also detected a significant fraction of cells in S-phase that had no transcripts.

If transcription is solely restricted to S phase, cells that arrest outside of S phase should produce no transcripts. To test this idea, we used an analog sensitive *cdc2-asM17* (K79E F84G) allele ^64^ which arrests cells before entry into S-phase (G1/S or G2/M transition) upon addition of 1NM-PP1. Upon 1NM-PP1 addition, we observed successful arrest since cells continue to elongate but fail to divide for up to 8 hours. We measured *dh* transcription at two-hour intervals and found that these arrested cells still produced transcripts (**Figure 3H**). We plotted the distribution and average numbers of transcripts per cell (0hr: 0.77 *dhF*, 1.52 *dhR* /cell; 8hr: 2.45 *dhF*, 4.67 *dhR* /cell) throughout the arrest period (**Supplementary Figure 3D-F**). These findings support the conclusion that pericentromeric transcription can occur outside S phase. One possibility is that low transcript counts bias our measurements. To raise baseline transcript levels without fully disrupting pericentromeric heterochromatin, we introduced a mutation in the Clr4 SET domain (R406H) which attenuates enzymatic activity but preserves silencing ^65^. This led to a 5-fold increase in average transcript counts compared to wild-type cells (**Supplementary Figure 3G,H**). Nevertheless, we observed no cell-cycle dependence in this background and no significant difference in transcript numbers across G2/M/G1 and S phase cells (**Supplementary Figure 3I-N**).

One other possibility is that transcription dynamics at pericentromeres could be fundamentally distinct from canonical transcription. A signature feature of RNA polymerase II transcriptional dynamics is that transcript abundance scales linearly as a function of cell volume, pointing to a homeostatic mechanism that maintains the overall RNA concentration in individual cells (**Figure 3I**) ^44,66,67^. If transcription from pericentromeric repeats is distinct from elsewhere in the genome, we predicted that the *dh* and *dg* transcript abundance would deviate from this homeostatic scaling model. In *clr4Δ* and in *ago1Δ* cells where transcripts are abundant, we observed that *dg* and *dh* transcripts scale linearly as a function of cell length much like all other transcripts in the *S*.*pombe* genome (**Figure 3J**). Even in the case of wild-type cells, where transcripts are rare, we observe the same linear scaling principle although the small number of transcripts produces large variance in our measurement. Lastly, *cdc2-asM17* cells which increase substantially in length also show scaling with longer cells producing more transcripts at 8hr compared to 0hr, which overall corresponds to a ∼3-fold increase in both *dhF* and *dhR* (**Supplementary Figure 3F)**. Based on these results, we concluded that the dynamics of RNA polymerase II transcription are no different at pericentromeric repeats compared to the rest of the genome.

### Modeling siRNA-directed methylation and gene silencing

Based on our data, we hypothesized that pericentromeric heterochromatin permits a low level of transcription throughout the cell cycle. To capture these dynamics, we developed a mathematical framework which assumes that transcript production is solely governed by feedback involving siRNA biogenesis and H3K9 methylation (**Figure 4A**). Our framework builds on important theoretical foundations developed earlier for RNAi and non-RNAi based heterochromatin systems ^68–71^, with two major changes: **(1)** low levels of small RNAs are produced in the absence of H3K9 methylation (priRNAs) ^72^ and **(2)** a nascent transcript is required for pericentromeric heterochromatin establishment ^29^ (see **Figure 4B** and **Supplementary Note 2** for additional details about the model).

**Figure 4.**
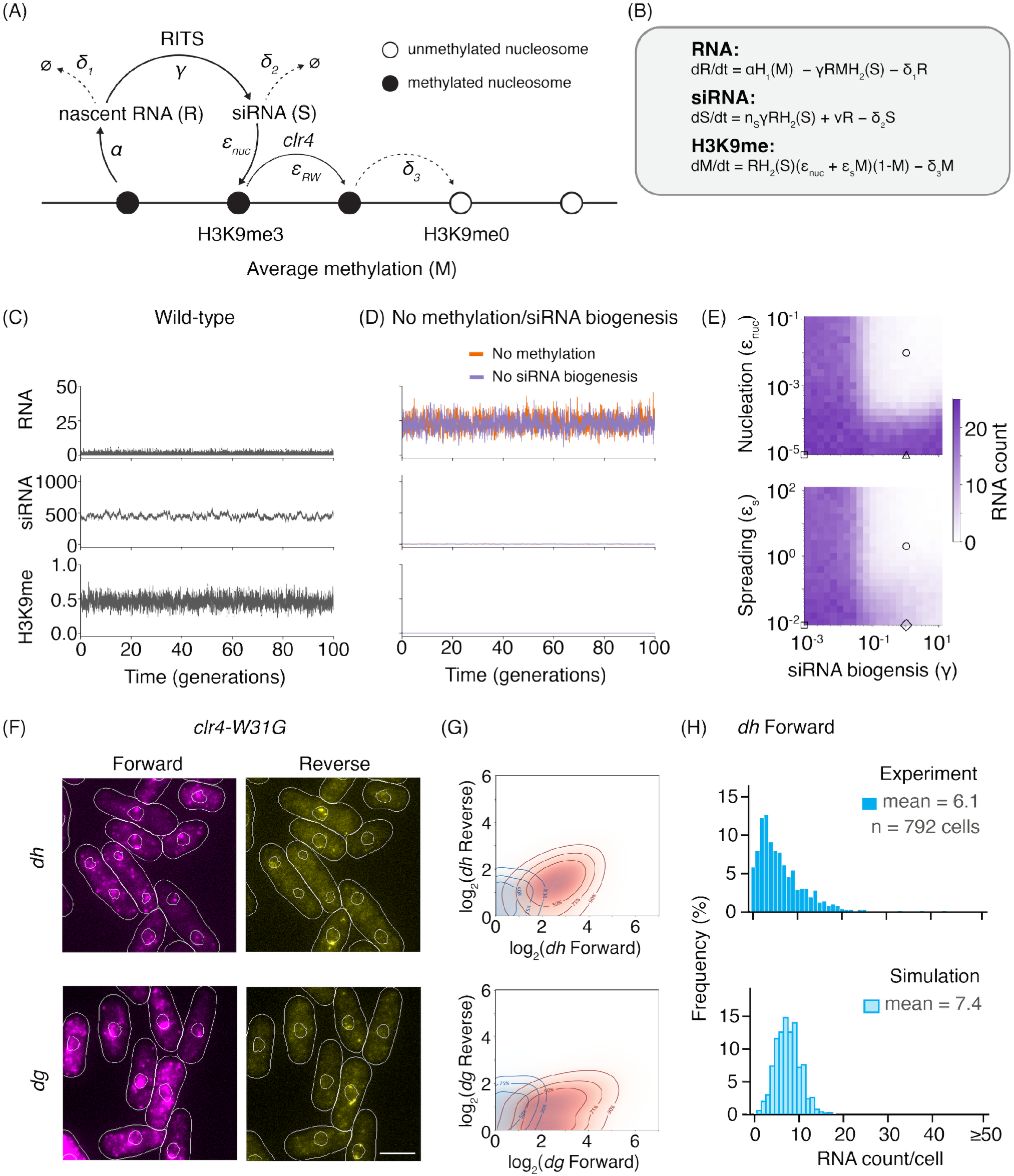
Modeling siRNA-directed methylation and gene silencing. **(A)** The *S*.*pombe* RNAi is modelled as a multi-variable system where the levels of RNA, H3K9me3 and siRNA mutually depend on each other. The nascent transcripts from partially methylated chromatin are targeted for processing and siRNA is generated. siRNAs recruit the RITS complex to the nascent transcripts which leads to reinforcement of H3K9me3. Here, α is the transcription rate, γ is the siRNA biogeneis rate, n_S_ is the number of siRNAs produced per co-transcriptional silencing event, ν is the primal siRNA production rate, ε_nuc_ and ε_s_ are the nucleation and spreading rates for methylation marks, δ_1_, δ_2_ and δ_3_ are the loss rates of RNA, siRNA and methylation marks. **(B)** The set of deterministic equations describing the coupling between the variables R, S and M in the model. **(C)** A stochastic trajectory simulated by Gillespie algorithm for wild-type cells show stable, low level transcription and moderate levels of H3K9me and siRNA. **(D)** A stochastic trajectory simulated with no methylation activity (ε_nuc_, ε_s_ = 0) or siRNA biogenesis (γ, ε_nuc_, ε_s_ = 0). This leads to complete loss of silencing, high transcription levels and absence of H3K9me and siRNA. **(E)** The phase diagrams show expected changes in transcript levels across a range of model parameters (γ, ε_nuc_ and ε_s_). The markers indicate the parameters used in the model to simulate wild-type (circle), *clr4Δ* (triangle), *ago1Δ* (square) and *clr4-W31G* (diamond). **(F)** Fluorescence images of *dg* and *dh* forward and reverse transcripts in *clr4-W31G* mutant. Scale bar is 5 μm in all the images. **(G)** The contour plots for *dg* and *dh* transcripts in *clr4-W31G* mutant are shown. Blue and red contours represent the wild-type and the mutant cells respectively. **(H)** Simulated RNA levels for *clr4-W31G* (ε_s_ = 0) agree well with the experimental data. The silencing is partially lost with RNA levels being higher than wild-type but lower than *clr4Δ*.

The deterministic equations assuming parameters consistent with experimental data reach a steady state in which non-coding RNA transcripts are expressed at low levels (∼1 per cell), while siRNA and H3K9me levels are constant at high levels (**Supplementary Figure 4A**). Stochastic simulations (Gillespie) show the trajectories of RNA, siRNA and H3K9me levels varying over time in a single cell (**Figure 4C**). Across 500 independent trajectories simulated over 100 generations, we observed no excursions into a high-expression state (**Supplementary Figure 4B**), indicating that silencing constrains transcription and this process is unperturbed by noise. Notably, the stationary distribution also predicts that a substantial fraction of cells (∼40%) contain no transcript at any given time matching our experimental data (**Supplementary Figure 4C**).

To test the model against known phenotypes, we set the methylation rate to zero to simulate a *clr4Δ* scenario. Transcripts stabilized at high levels (∼20 per cell) while siRNA and H3K9me levels completely collapsed which is reflected both in our stochastic simulations and deterministic equations (**Figure 4D** and **Supplementary Figure 4D-F**). Setting siRNA biogenesis rates to zero to simulate RNAi pathway mutants essentially produced the same outcome (**Figure 4D** and **Supplementary Figure 4G-I**). Both results recapitulate the near-identical loss of silencing phenotypes we experimentally measured by smFISH in *clr4Δ* and *ago1Δ* cells (**Figure 2** and **Supplementary Figure 4F,I**).

H3K9 methylation spreading involves Clr4 ‘read-write’ activity and a mutation within the Clr4 chromodomain (*clr4-W31G*) disrupts its ability to bind to H3K9me ^31,73^. smFISH measurements in *clr4-W31G* cells show *dg* and *dh* forward and reverse transcripts increased ∼10-fold relative to wild-type cells (**Figure 4F-H**). This increase in transcript count was still lower than what we observe in RNAi mutants or *clr4Δ* cells. In simulations where we disrupt “read-write” i.e. spreading from existing methylation is set to zero (ε_s_ = 0), transcripts were also expressed at higher levels quantitatively matching our experimental data (**Figure 4H**). Hence, our model captures how different aspects of feedback strength (siRNA and H3K9me) within the context of RNAi mediated heterochromatin establishment autonomously regulate transcript production without any dedicated mechanism that licenses their control.

### Tuning demethylase activity defines distinct regimes for silencing and switching

A second orthogonal parameter in our model that affects H3K9me abundance is the rate of H3K9 removal (δ_3_) (**Figure 4A**). The most critical factor that promotes H3K9me erasure in *S*.*pombe* is the putative H3K9me demethylase, Epe1 ^74–77^. In our simulations, increasing Epe1 activity relative to wild-type cells leads to a monotonic decrease in H3K9me and a concomitant increase in transcription (**Figure 5A,B**). siRNA levels initially increase but we reach a tipping point when demethylation rates are an order of magnitude higher than in wild-type cells when siRNA levels start to decrease converging on levels that we observe in *clr4Δ* cells (**Figure 5C**). This is also reflected in our stochastic simulations in an Epe1 overexpression scenario that produces stable, yet high levels of transcripts (**Figure 5D**). To test these predictions, we overexpressed Epe1 using a thiamine repressible *nmt41* promoter. In no thiamine media, Epe1 overexpression results in a substantial increase in transcript production (>90%) compared to wild-type cells (50%) (**Figure 5E,F** and **Supplementary Figure 5A,B**) ^77^. The average number of *dg* and *dh* transcripts in all directions (*dhF*: 13, *dhR*: 5, *dgF*: 12, *dgR*: 6 RNA/cell) is much higher than in a wild-type background.

**Figure 5.**
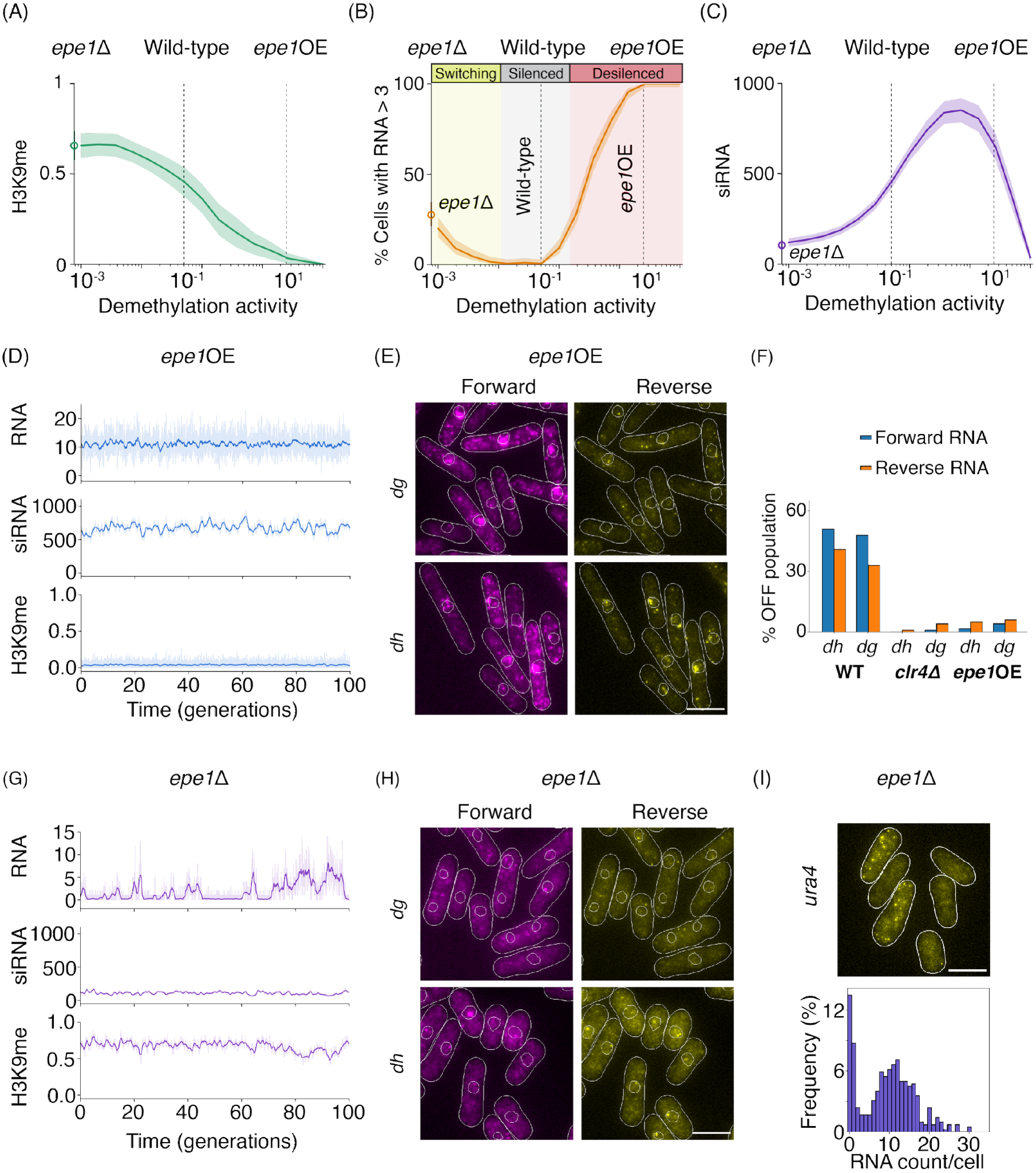
Tuning demethylase activity defines distinct regimes for silencing and switching. **(A)** Methylation declines with increase in demethylation activity. The mean values and the spread are obtained from 200 simulations for each parameter choice. **(B)** Percentage of cells with RNA > 3 increases when the demethylation activity either increased or decreased relative to wild-type. **(C)** siRNA levels rise with increasing demethylation activity, then collapse at higher demethylation, reflecting loss of RNAi. **(D)** Simulation traces for RNA, siRNA and H3K9me when Epe1 is overexpressed (*epe1*OE). **(E)** Fluorescence images of *dg* and *dh* transcripts in *epe1*OE conditions. Scale bar is 5 μm in all the images. **(F)** Percentage of cells with no RNA is quantified for *dg* and *dh* transcripts in various backgrounds. **(G)** Simulation traces for RNA, siRNA and H3K9me for *epe1Δ*. **(H)** Fluorescence images for *dg* and *dh* transcripts in *epe1Δ*. **(I)** Top panel shows *ura4+* transcripts imaged in *epe1Δ* and the bottom panel shows the histogram for RNA counts per cell.

Decreasing the demethylation rate increases H3K9me levels, but transcription rises abruptly when H3K9me levels fall below those in wild-type cells (**Figure 5A,B**). The stochastic simulations show large fluctuations relative to the silenced state resulting in switching between high and low transcript numbers (**Figure 5G**). We confirmed that these state changes do not reflect bistability since the deterministic solutions of our model predicts a single fixed point with high H3K9me levels and low levels of transcripts compared to wild-type cells (**Supplementary Figure 5D**). In our experiments, deleting Epe1 supports the stochastic switching model since both *dg* and *dh* expressing cells split into two populations, cells with transcripts (*dhF*: 32%, *dhR*: 45%, *dgF*: 22% *dgR*: 59%) and cells without transcripts (*dhF*: 68%, *dhR*: 55%, *dgF*: 78% *dgR*: 41%) (**Figure 5H** and **Supplementary Figure 5F,G**). The distinction in transcript levels between the two populations is even more striking for *ura4+* (**Figure 5I** and **Supplementary Figure 5H**). This may be because *ura4+* is a single copy reporter within pericentromeric repeats as opposed to *dg* and *dh* which are repeats with each copy being able to switch independently. In cells that express *ura4+*, transcript abundance matches what we typically observe in a *clr4Δ* background (70%) whereas the rest of the population was wild-type like with low transcript numbers (∼30%). Hence, these results suggest that enhancing silencing disrupts feedback leading to stochastic switching.

### Real-time single cell imaging reveals that switching is a dynamic feature in *epe1Δ* cells

To determine whether cells dynamically switch between expression states as predicted by our model, we replaced the *ura4+* reporter with a *GFP* reporter to measure real-time changes in gene expression. As expected, wild-type cells have low GFP expression and *clr4Δ* cells have high GFP expression demonstrating that our reporter is sensitive to the normal genetic requirements of H3K9 methylation and heterochromatin establishment (**Figure 6A, Supplementary Figure S6A**). However, across three independent *epe1Δ* biological replicates, we observed a bimodal population of cells where 57% of cells are in the ON state and 43% (mean, n = 3 replicates) of cells are in the OFF state (**Figure 6A** and **Supplementary Figures S6B,C**). To determine whether these are stable or dynamic populations, we continuously imaged individual cell lineages using a custom microfluidic device for 4 days (∼ 40 cell generations) as shown in **Figure 6B**. About ∼60% of individual *epe1Δ* cell lineages stochastically switch between ON and OFF expression states (**Figure 6C**). Hence, switching represents *epe1Δ* cells dynamically switching between distinct transcriptional states. Although ∼40% of cells did not exhibit switching on the timescale of our measurement, we anticipate that this is likely because the switching rate is heterogenous and slow switching lineages are likely never captured during the time-course of our measurement (4 days).

**Figure 6.**
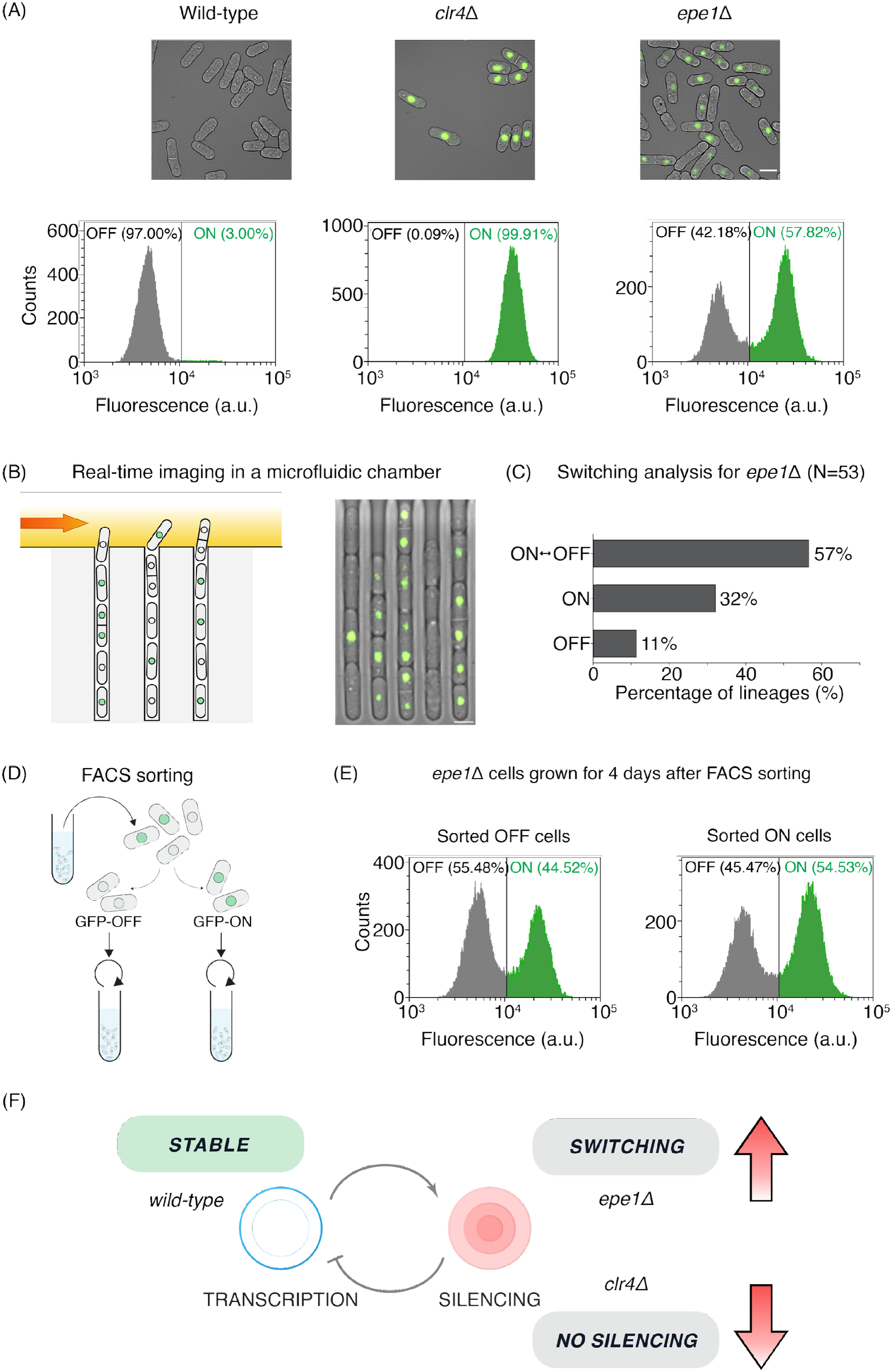
Real-time single cell imaging reveals that switching is a dynamic feature in *epe1Δ* cells. **(A)** Fluorescence images of centromeric GFP reporter in wild-type, *clr4Δ* and *epe1Δ*. Corresponding FACS analysis shows unimodal distributions for wild-type and *clr4Δ* and bimodal for *epe1Δ*. Scale bar is 5 μm in all the images. **(B)** Real-time imaging of *epe1Δ* cells loaded in the microfluidic chamber.**(C)** Quantification of single-cell switching behavior for a pericentromeric GFP reporter (*otr1R::mNeonGreen-sfGFP*). **(D)** FACS sorting was performed on *epe1Δ* cells to isolate ON and OFF cells. **(E)** The isolated cells were regrown separately for 4 days upon which the FACS analysis was done. Both the sorted ON and OFF populations reach an equilibrium with a mixed population. **(F)** Summary model: Stable heterochromatin maintenance in RNA-based silencing systems depends on balancing methylation and transcription levels.

We sorted the top 25% of ON or OFF cells based on GFP expression levels (**Figure 6D**) from three independent biological replicates. If cells dynamically switch between expression states in an *epe1Δ* background, we expect that our sorted GFP-ON and GFP-OFF populations will re-equilibrate and achieve a new steady state containing a mixture of GFP-ON and GFP-OFF cells. Indeed, after ∼ 4 days of growth in liquid culture, both sorted populations reached the same steady state consisting of a mixture of ON and OFF cells (**Figure 6E** and **Supplementary Figures S6B,C**). These results suggest that enhancing silencing within pericentromeric repeats by deleting Epe1 leads to cells dynamically switching between distinct expression states. These observations are consistent with previous studies where deleting Epe1 produces variegated phenotypes in plate based assays ^78,79^.

## Discussion

The widespread adoption of RNA-based strategies to recruit silencing factors poses a fundamental paradox: how do two antagonistic processes such as transcription and silencing coexist to regulate gene expression? ^37^ We resolve this paradox by demonstrating that low-levels of non-coding transcription produces nascent RNA scaffolds that promote robust heterochromatin establishment independent of any inputs from the cell-cycle. Instead, our model suggests that transcription dynamics are autonomously regulated based on feedback involving H3K9 methylation and siRNA biogenesis. Using single-molecule FISH (smFISH), we achieved selection- and enrichment-independent measurements of pericentromeric transcripts in unperturbed *S*.*pombe* cells (**Figure 2**). Although smFISH is a fixation-based approach, our measurements can directly measure endogenous RNA molecules without additional modifications to the transcript or locus ^43,80,81^.

Our selection and enrichment independent approach enabled us to detect rare transcripts across all phases of the cell-cycle. All classes of pericentromeric transcripts independent of orientation were detected at any point with no bias in expression. These observations underscore the stochastic and heterogenous nature of pericentromeric transcription. The basal production of pericentromeric transcripts across multiple repeats ensures that nascent RNA scaffolds are continuously available for the RITS complex to bind and re-establish heterochromatin ^29^.

Although the exact molecular mechanism for what causes transcript production is not specified by our model, one plausible candidate is a stochastic change in H3K9 methylation valency through enzyme activity (H3K9me2 versus H3K9me3) or nucleosome turnover ^82^. Previous studies have shown that siRNA biogenesis remains intact when Clr4 can catalyze H3K9me2 but not H3K9me3, demonstrating that H3K9me2 can support a transcriptionally permissive state ^83^. A second possible mechanism is that transcription factors such as the PhpC^NF-Y^ complex can bind to pericentromeric heterochromatin independently of siRNA to activate transcription ^84^. Importantly, our proposed framework need not invoke any explicit mode of regulation. In principle, even baseline levels of RNA polymerase II transcription within pericentromeric repeats can produce rare transcripts to fuel heterochromatin establishment ^47,48^.

Based on measuring strand-specific non-coding RNA transcription, we find a striking pattern of nuclear localization for *dgR* and *dhR* transcripts. This is in contrast to other non-coding transcripts such as *dgF* and *dh*F which are successfully exported to the cytoplasm (**Figure 2H)**. It is possible that sequence-intrinsic features corresponding to *dgR* and *dhR* such as an RNA secondary structure or cryptic introns may control nuclear retention ^85–88^. Notably, *dhR* when expressed from an ectopic site is a strong nucleator of RNAi mediated heterochromatin formation ^46^ and other studies have shown that, in the absence of RNAi, the MTREC/PAXT complex binds *SPNCRNA*.*230*, which contains the *dhR* repeat ^89^. These observations reveal how cells might exploit sequence-intrinsic features of non-coding RNA transcripts to nucleate heterochromatin.

We propose that Epe1 activity is the primary axis that determines the steady-state transcriptional set point in wild-type cells since altering its function forces cells to adopt distinct transcriptional profiles. Epe1 is a putative H3K9 demethylase that antagonizes heterochromatin assembly ^74–77^ . Yet, its deletion simultaneously destabilizes and promotes heterochromatin establishment within pericentromeric repeats ^78,79^. To reconcile these conflicting phenotypes, previous studies proposed non-canonical functions for Epe1, such as direct transcriptional activation or histone acetyltransferase recruitment ^77,90^. Instead, our simulations and experimental data show stochastic switching upon deleting Epe1 is a direct consequence of the feedback architecture. In *epe1Δ* cells, unchecked H3K9 methylation leads to enhanced silencing within pericentromeric heterochromatin which in turn completely eliminates transcript production. In the absence of a nascent RNA scaffold to recruit silencing complexes and sustain siRNA biogenesis, the locus switches into a fully expressed state. Low levels of priRNAs are then required to re-nucleate heterochromatin which accounts for how *epe1Δ* cells stochastically switch between long-lived silent and expressed states that are stable across multiple generations.

Based on our findings, we propose that RNA-based silencing architectures may provide a tunable mechanism for establishing and maintaining stable transcriptional set points without the explicit need for external regulatory inputs. In *S*.*pombe*, the RNAi feedback architecture involving siRNA biogenesis, and H3K9 methylation establishes a set point where transcription produces an average of one to two transcripts per cell. The evolution of different modes of feed-back control is also apparent if we look beyond *S*.*pombe*. For example, fungi such as *Mucorales* lack Chp1, a key adapter protein which in *S*.*pombe* couples H3K9me establishment to siRNA biogenesis ^91^. Several fungal lineages lack the Argonaute chaperone complex proteins Arb1 and Arb2, which are important for siRNA loading and stability in *S*.*pombe* ^91,92^. Additionally, RNA-directed DNA methylation in plants and Piwi-interacting RNA-mediated silencing in animals represent distinct evolutionary manifestations of RNA-guided epigenetic control ^18–21^. In each instance, we propose that these systems have adopted distinct feedback architectures which defines their unique transcriptional set points. Hence, we propose that transcriptional regulation can emerge from feedback without requiring dedicated signaling pathways, providing flexibility in how RNA-based silencing systems evolve to control transcription.

## Acknowledgements

We thank members of the Ragunathan lab for their insightful feedback in helping edit the manuscript and interpret the data. We thank Neha Arora, Emily Metcalf and the Whitehead Institute Flow Cytometry Core facility for help with cell sorting. We thank Juan Serrano Jimenez and Iris Nava for help with building reporter strains for pericentromeric silencing assays. We thank Hiten Madhani, Danesh Moazed, and Songtao Jia for sharing fission yeast strains used in this study. This work was supported by an NIH/NIGMS Award (R35GM137832) to K.R., an American Cancer Society Award (RSG-22-117-01-DMC) to K.R., NSF CAREER Award (2441697) to K.R. and Human Frontier Science Program Fellowship (LT0020/2024-L) to A.Z.A. This paper was typeset with the bioRxiv word template by @Chrelli: www.github.com/chrelli/bioRxiv-word-template.

## Author contributions

S.K.M. conceptualized the project, developed the methodology, performed the investigation, analyzed the data, visualized the data, and reviewed/edited the final manuscript, A.Z.A. performed the investigation, analyzed the data, and reviewed/edited the final manuscript, S.C. performed the investigation, analyzed the data, visualized the data, and reviewed/edited the final manuscript, D.M. performed the investigation and analyzed the data, and reviewed/edited the final manuscript, J.K. developed the methodology, visualized the data, and reviewed/edited the final manuscript, K.R. conceptualized the project, analyzed the data, visualized the data, wrote the original draft, reviewed/edited the final manuscript, and supervised the study,

## Materials and Methods

### Plasmids

Plasmid expressing eGFP-PCNA was constructed using Gibson assembly. The *eGFP-pcn1* cassette including the promoter and terminator was PCR amplified from a previously published plasmid construct (Addgene pAV0620)^93^. The PCR product is assembled into an integration vector with homology to *lys3* locus for ectopic insertion into *S*.*pombe* genome.

### Strains

The *S*.*pombe* strains that carried *ura4-DS/E* at the endogenous *ura4* locus were transformed with a synthetic gene block containing the blasticidin (BSD) resistance marker. The strains carrying the *clr4* point mutants R406H and W31G were generated by CRISPR-Cas9 targeting the endogenous *clr4* locus, followed by repair with a synthetic gene block encoding the respective mutations. Endogenous Cnp1 was tagged at its C-terminus with an sf-GFP fusion construct by transformation of a cassette generated by overlap extension PCR, which has the homology to sequences flanking the endogenous stop codon. Centromeric fluorescent reporter (*otr1R::mNeonGreen-msfGFP*) was constructed by replacing *ura4+* at the *otr1R* locus of *cen1* with a *pcam1-mNeonGreen-msfGFP-Ura4t* cassette into *epe1Δ* and *clr4Δ* backgrounds. The cassette was generated by overlap extension PCR with homology to the *ura4* locus and integrated by CRISPR-Cas9 targeting the *ura4* locus. Wild-type strain carrying this fluorescent reporter was obtained by crossing the *clr4Δ* reporter strain with a wild-type stain lacking the reporter. All the yeast transformations were performed by electroporation method. For all the experiments, cells were grown to mid-log phase (OD600 0.5 - 0.8) in YEA media at 32ºC, unless stated otherwise. To arrest *cdc2-asM17* cells, 1NM-PP1 was added to the culture medium at a final concentration of 2 µM. Strains used in this study are listed in **Supplementary Table 1**.

### Single-molecule RNA-FISH

The sample preparation protocol was adapted from earlier reports ^43,94,95^. The cells were fixed using 3% formaldehyde at room temperature on a rotator for 20 min and then spheroplasted using 2mg/ml Zymolyase at 37ºC for upto 1hr. Spheroplasting was monitored by microscopy after adding 2% SDS and was considered complete when all the cells appeared translucent. Cells are fixed in 70% ethanol at room temperature for 4 hrs and incubated in Formamide Wash buffer (FWB: 10% Formamide in 2x SSC) for 30 min at room temperature. Hybridization reaction is setup with FISH probes (Stellaris Biosciences, see **Supplementary Note 1 for probe s**equences) at a final concentration of 25 nM overnight in FWB at 30ºC with shaking at 75 rpm. Cells are then resuspended in FWB and incubated at 30ºC for 30 min, and the cells are washed once again. Cells were resuspended in FWB containing DAPI (1µg/ml final concentration). Cells are washed in FWB and resuspended in imaging buffer (2x SSC, 50 mM at Tris pH8). Samples were stored at 4ºC until they are imaged.

Fluorescence images were acquired using a widefield epifluorescence microscope equipped with a 100×/1.49 NA oil immersion objective and DAPI/GFP/Cy3/Cy5 filter sets. Exposure times and acquisition settings were kept constant for all samples. Image analysis was performed in Python using Cellpose ^96,97^ for cell segmentation by training a model with a dataset of manually labelled dataset of cells. Elongated *cdc2-asM17* cells arrested with 1-NM-PP1 were segmented manually. Cell length was estimated as the major axis of the ellipse fitted to each segmented cell. Nuclear segmentation and spot analysis were performed using customized scripts built on Big-FISH ^98^. smFISH images were analyzed on 2D maximum-intensity projections of the z-stacks. For each dataset, the intensity threshold and spot size parameters for spot detection were chosen by visually inspecting detected spots in representative images and kept constant across all images in that dataset.

To visualize forward and reverse transcript counts together, contour plots were generated by estimating their joint distribution using a two-dimensional Gaussian kernel density with the bandwidth set to twice Scott’s rule for smoothing. Counts were transformed as log_2_ (1+RNA/cell) before density estimation. Contour levels were set from the kernel density values at individual cells, so that the 50%, 75% and 90% contours enclose approximately those fractions of cells in each sample.

### Microfluidics

For single-cell time-lapse imaging, cells were loaded into a custom-fabricated PDMS-based microfluidics device. Fresh YEA media was perfused continuously at a flow rate of 40 µL/min, and the device was maintained at 32°C throughout the experiment. Cells were imaged for 4 days with snapshots of GFP emission captured every 30 min to limit phototoxicity. Images were acquired using a widefield epifluorescence microscope equipped with a 20× objective and a GFP filter set. Image processing was done in Fiji (ImageJ) and a Gaussian filter was applied for displaying the images.

### Flow cytometry and cell sorting

Cells (1 OD) were resuspended in 1 ml YEA media and GFP fluorescence was measured on a CytoFLEX 4 flow cytometer (Beckman Coulter) using 488 nm excitation wavelength. Cells were gated on forward and side scatter to exclude debris and 20,000 cells were analyzed for each sample. Cell sorting was performed on a FACS Aria III (BD Biosciences) using 488 nm excitation wavelength. GFP expression and non-expressing cells were collected, with at least 500,000 live cells sorted for each sample.

### Gillespie simulations

The model was simulated with the stochastic simulation algorithm using GillesPy2 package ^99^ and custom Python code. For phase diagrams, each cell was simulated for 40 generations and RNA counts were recorded in 500 cells. Time-courses were recorded for 100 generations after a 20-generation wait time to let the system reach the equilibrium. See **Supplementary Note 2** for details.

