## Supplementary Figures for "Feedback control via H3K9 methylation and siRNA biogenesis autonomously maintains RNAi heterochromatin through continuous low-level transcription"

Supplementary Figure 1: Workflow for single-cell RNA quantification using smFISH

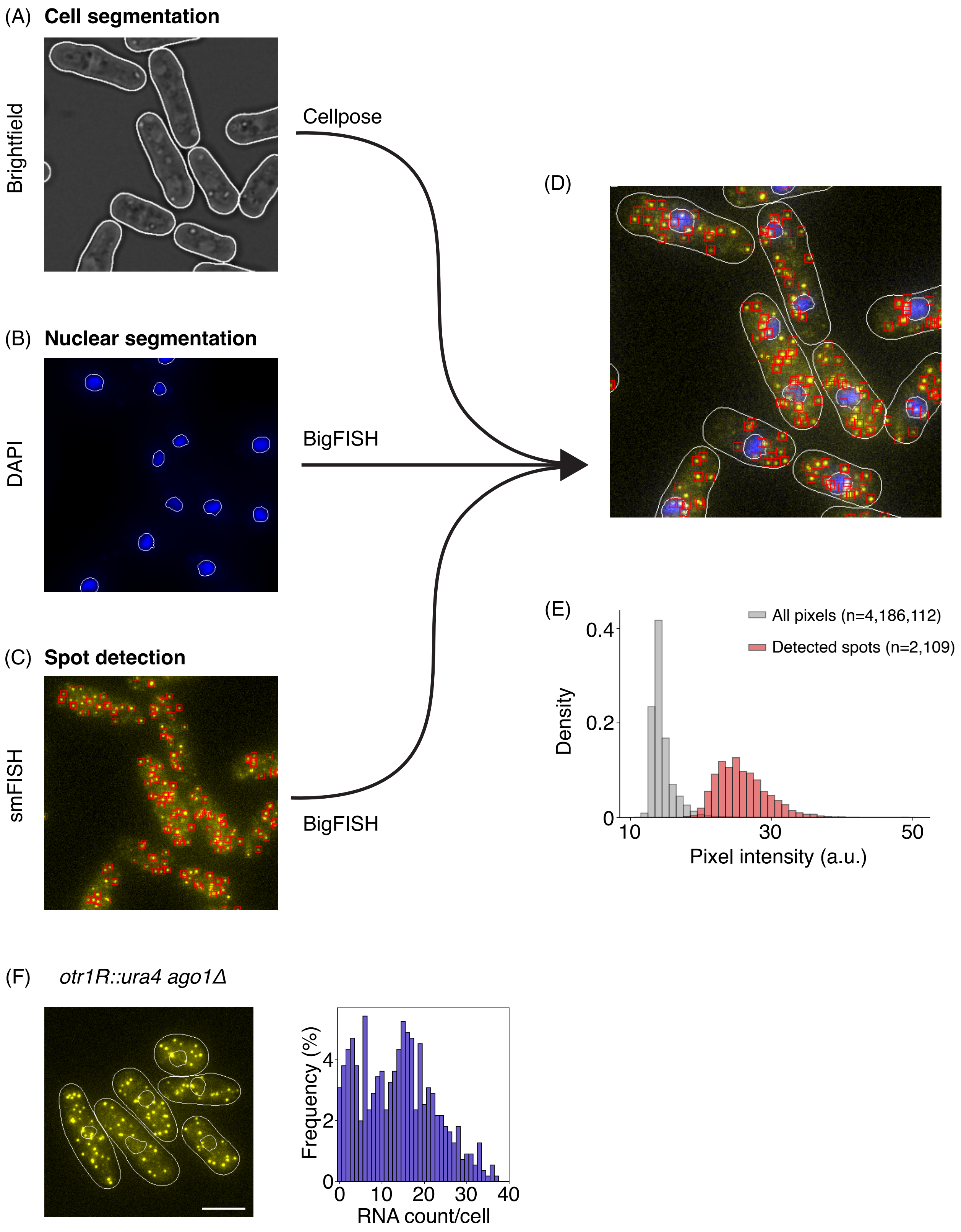

**Supplementary Figure 1: Workflow for single-cell RNA quantification using smFISH**

**(A)** Cell segmentation of brightfield images was performed using Cellpose. The detected cell boundaries are overlaid as white outlines.

**(B)** Nuclear segmentation of DAPI images was performed using BigFISH. The detected nuclear boundaries are overlaid as white outlines.

**(C)** Spot detection in smFISH images was performed using BigFISH. Red squares mark the detected spots.

**(D)** Combined output of cell segmentation, nuclear segmentation, and spot detection from the sequential workflow.

**(E)** Histograms comparing the intensities of detected spots with those of all pixels in the image.

**(F)** Fluorescence microscopy image of *ura4+* transcripts in *ago1Δ* cells. Histogram of RNA count per cell is shown to the right. Scale bar is 5 μm.

Supplementary Figure 2: smFISH controls and increased bidirectional pericentromeric transcription in heterochromatin pathway mutants

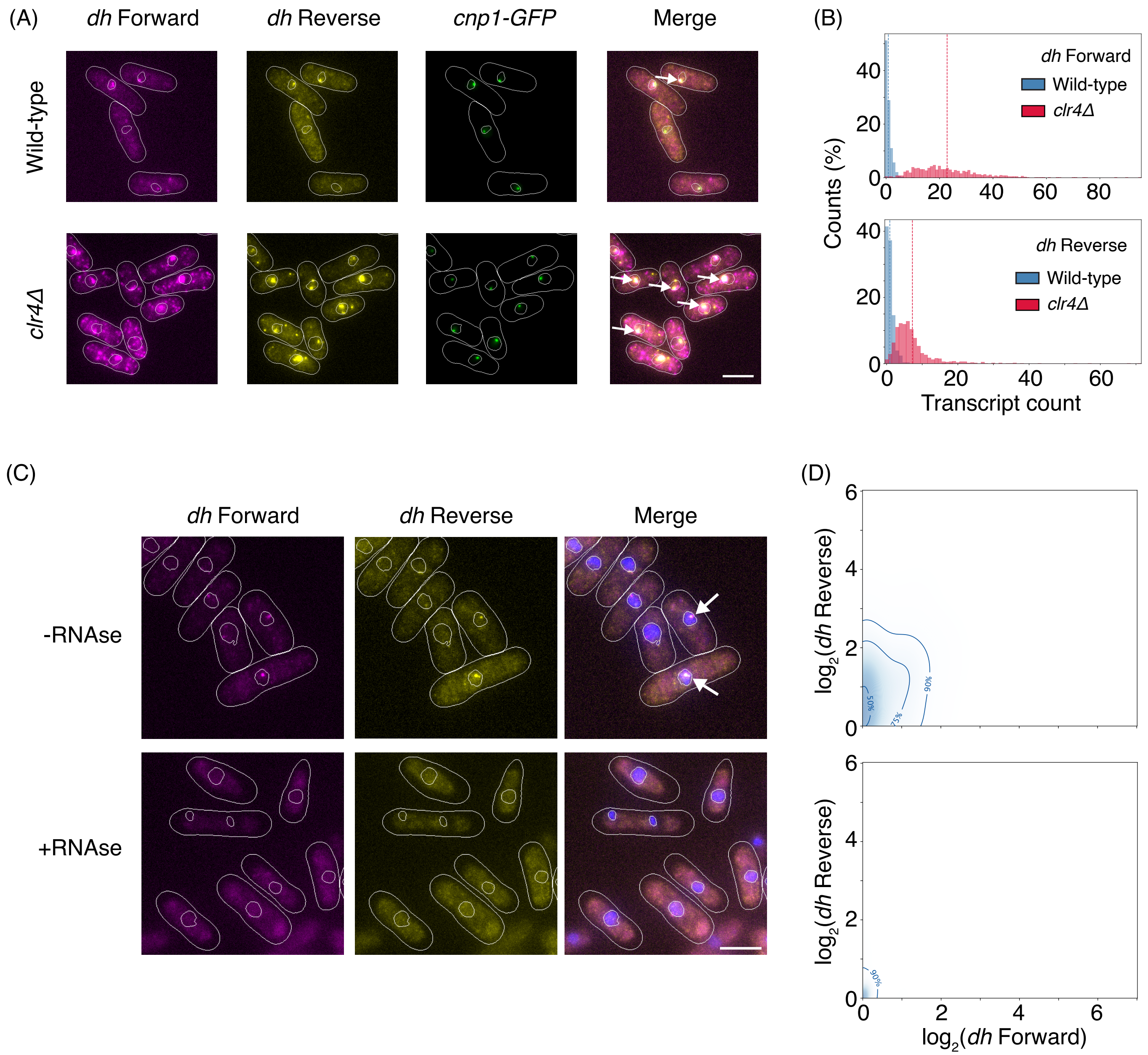

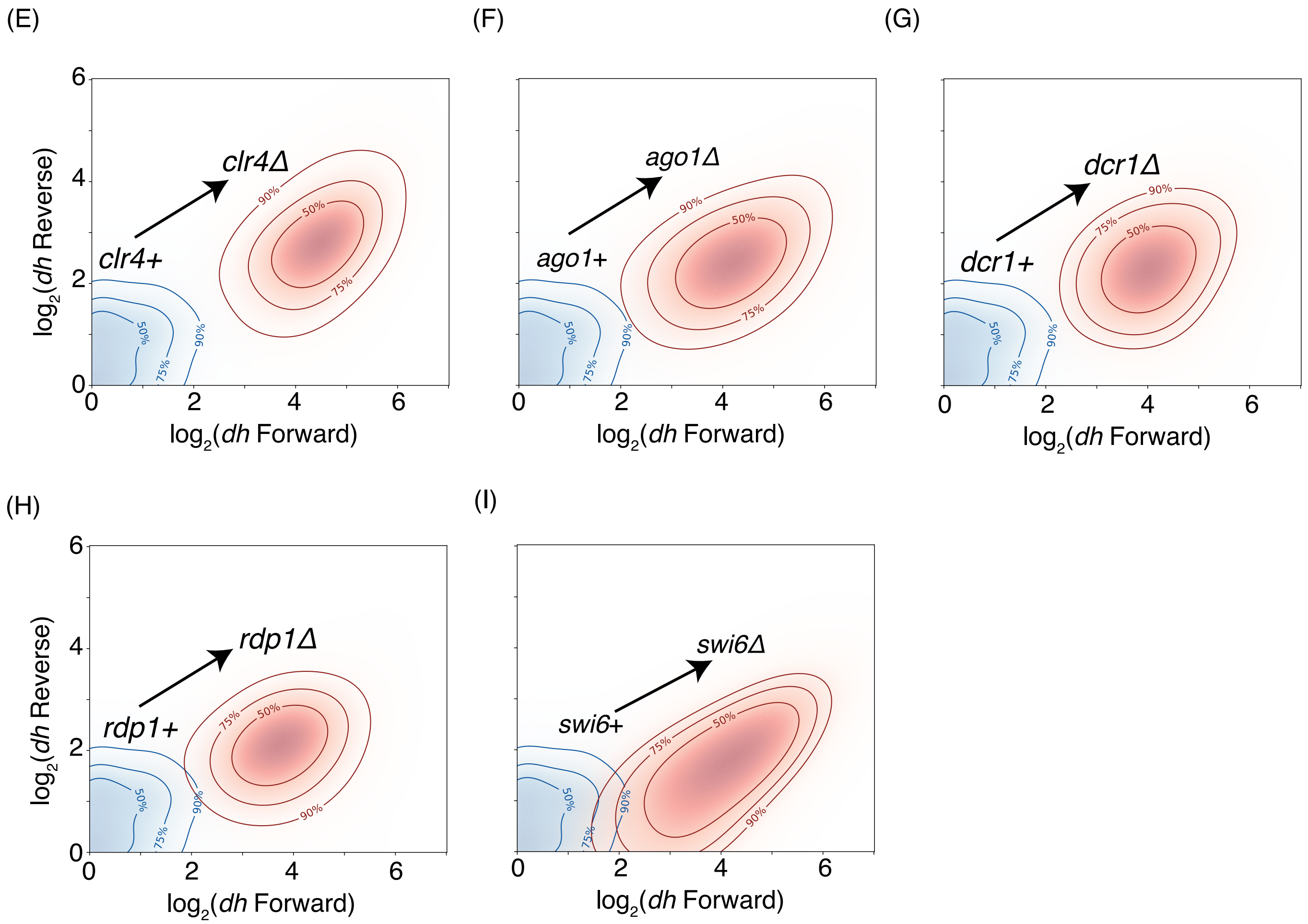
(E)
(F)
(G)
(H)
(I)
log2(dh Reverse)
log2(dh Forward)
clr4Δ
clr4+
ago1Δ
ago1+
dcr1Δ
dcr1+
rdp1Δ
rdp1+
swi6Δ
swi6+
90%
75%
50%
0
2
4
6

**Supplementary Figure 2: smFISH controls and increased bidirectional pericentromeric transcription in heterochromatin pathway mutants**

**(A)** Fluorescence images of *dh* forward (magenta) and reverse (yellow) transcripts in wild-type cells (top) and *clr4Δ* cells (bottom). Colocalization of transcripts with Cnp1-GFP confirms they originate from the centromere. Scale bar is 5 μm in all the images.

**(B)** Histograms of *dh* forward and reverse RNA counts per cell for wild-type (top) and *clr4Δ* cells (bottom).

**(C)** The *dh* forward and reverse transcripts are imaged before (top) and after (bottom) the RNAse treatment.

**(D)** The contour plots for *dh* forward and reverse transcripts before (top) and after (bottom) RNAse treatment.

**(E)** The contour plots for *dh* forward and reverse transcripts show increased pericentromeric transcription in both directions in *clr4Δ* cells. Blue and red contours represent the wild-type and the mutant cells respectively.

(**F-I**) Similar contour plots as (E) for *ago1Δ* (F), *swi6Δ* (G), *rdp1Δ* (H) and *dcr1Δ* (I).

Supplementary Figure 3: Cell cycle independent transcription of *dg* and *dh* repeats in wild-type and *clr4 R406H* cells

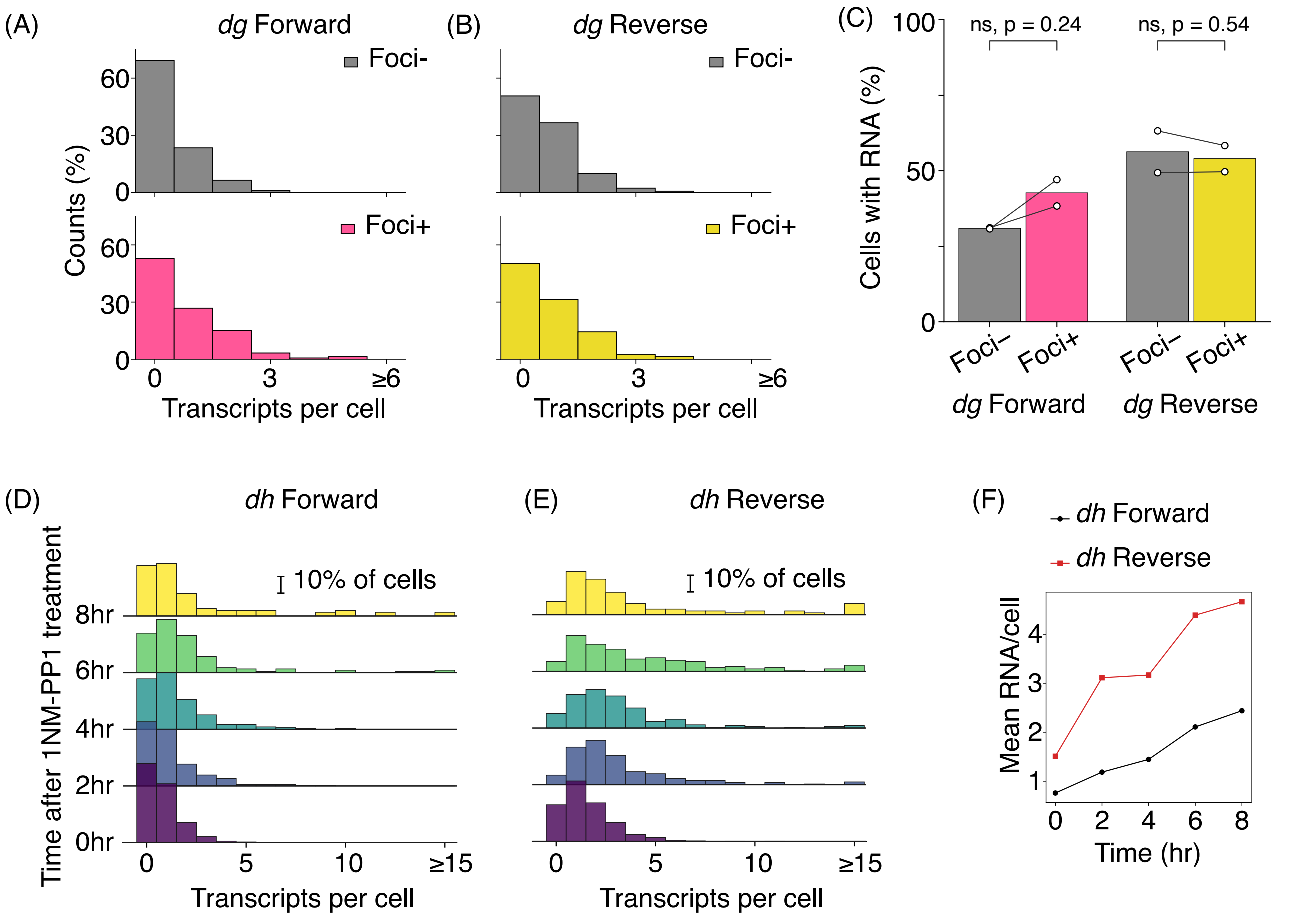

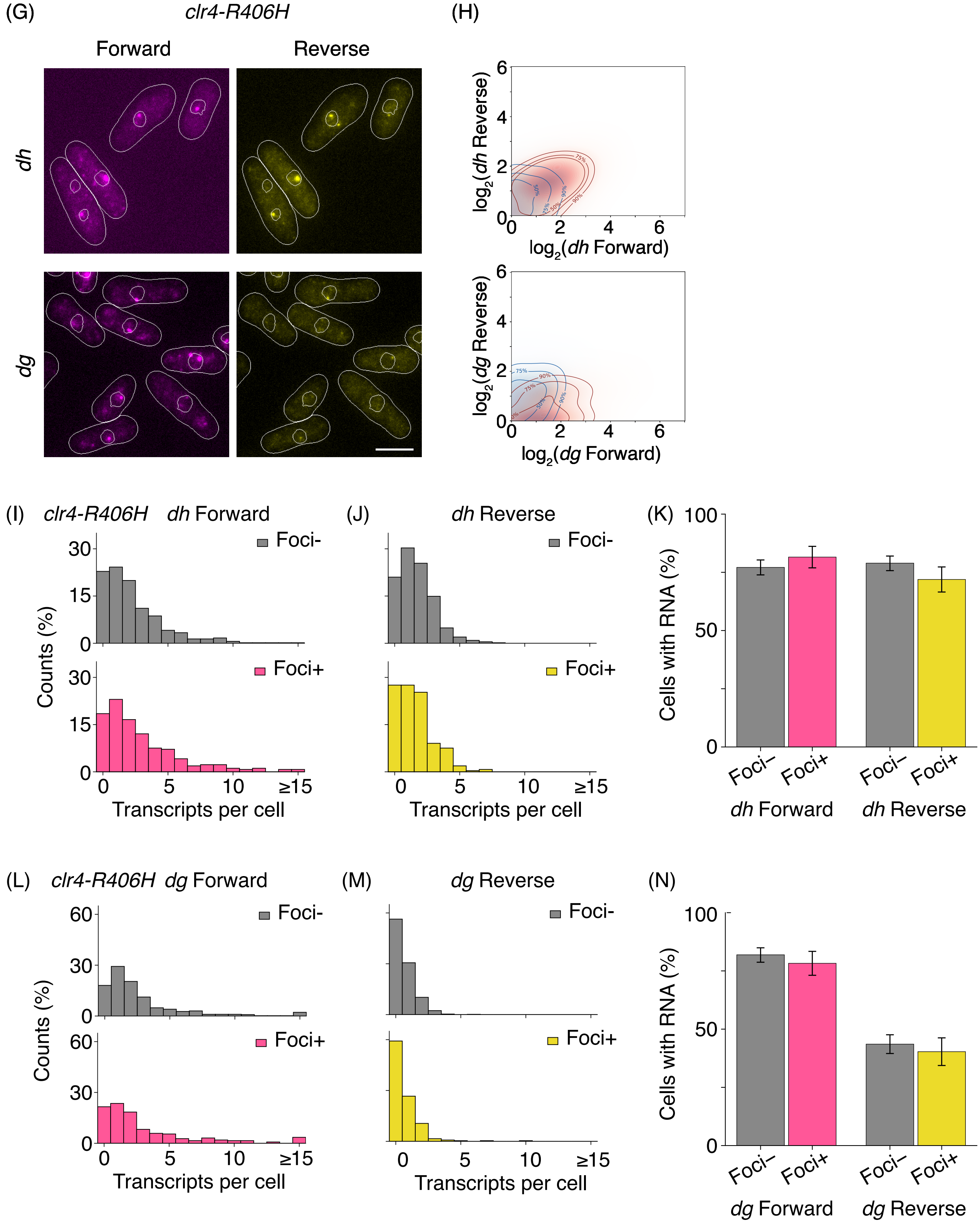
(G)
clr4-R406H
Forward
Reverse
dh
dg
(H)
log2(dh Reverse)
log2(dh Forward)
log2(dg Reverse)
log2(dg Forward)
0
2
4
6
(I)
clr4-R406H dh Forward
Foci-
Foci+
Counts (%)
Transcripts per cell
0
15
30
5
10
≥15
(J)
dh Reverse
Foci-
Foci+
Transcripts per cell
(K)
Cells with RNA (%)
0
50
100
Foci–
Foci+
dh Forward
dh Reverse
(L)
clr4-R406H dg Forward
Foci-
Foci+
Counts (%)
60
Transcripts per cell
(M)
dg Reverse
Foci-
Foci+
Transcripts per cell
(N)
Cells with RNA (%)
Foci–
Foci+
dg Forward
dg Reverse

**Supplementary Figure 3: Cell cycle independent transcription of *dg* and *dh* repeats in wild-type and *clr4-R406H* cells**

**(A)** Histograms of *dg* forward transcript numbers per cell in foci- and foci+ wild-type cells.

**(B)** Similar to (A) for *dg* reverse transcripts.

**(C)** Percentage of wild-type cells with *dg* RNA in foci- and foci+ cells. Data are from two replicates.

**(D)** Histograms of *dh* forward transcripts per cell at the indicated time points upon adding 1NM-PP1 to *cdc2-asM17* cells.

**(E)** Similar to (D) for *dh* reverse transcripts.

**(F)** Mean *dh* RNA per cell plot at indicated time points after 1NM-PP1-induced arrest.

**(G)** RNA-FISH images of *dg* and *dh* transcripts in *clr4-R406H* cells. Scale bar is 5 μm in all the images.

**(H)** Contour plots for *dg* and *dh* transcript levels in *clr4-R406H* cells. Blue and red contours represent the wild-type and the mutant cells respectively.

**(I)** Histograms of *dh* forward transcript numbers in cells *clr4-R406H* foci- and foci+ cells.

**(J)** Similar to (I) for *dh* reverse transcript.

**(K)** Percentage of *clr4-R406H* cells with *dh* RNA in foci- and foci+ cells. The error bars show 95% confidence intervals obtained by bootstrapping cells.

**(L)** Histograms of *dg* transcript numbers in *clr4-R406H* cells foci- and foci+.

**(M)** Similar to (L) for *dg* reverse.

**(N)** Percentage of *clr4-R406H* cells with *dg* RNA in cells with and without PCNA-GFP foci. The error bars show 95% confidence intervals obtained by bootstrapping cells. Scale bar is 5 μm in all the images.

Supplementary Figure 4: Simulated and experimental transcript abundance histograms for wild-type and mutant cells

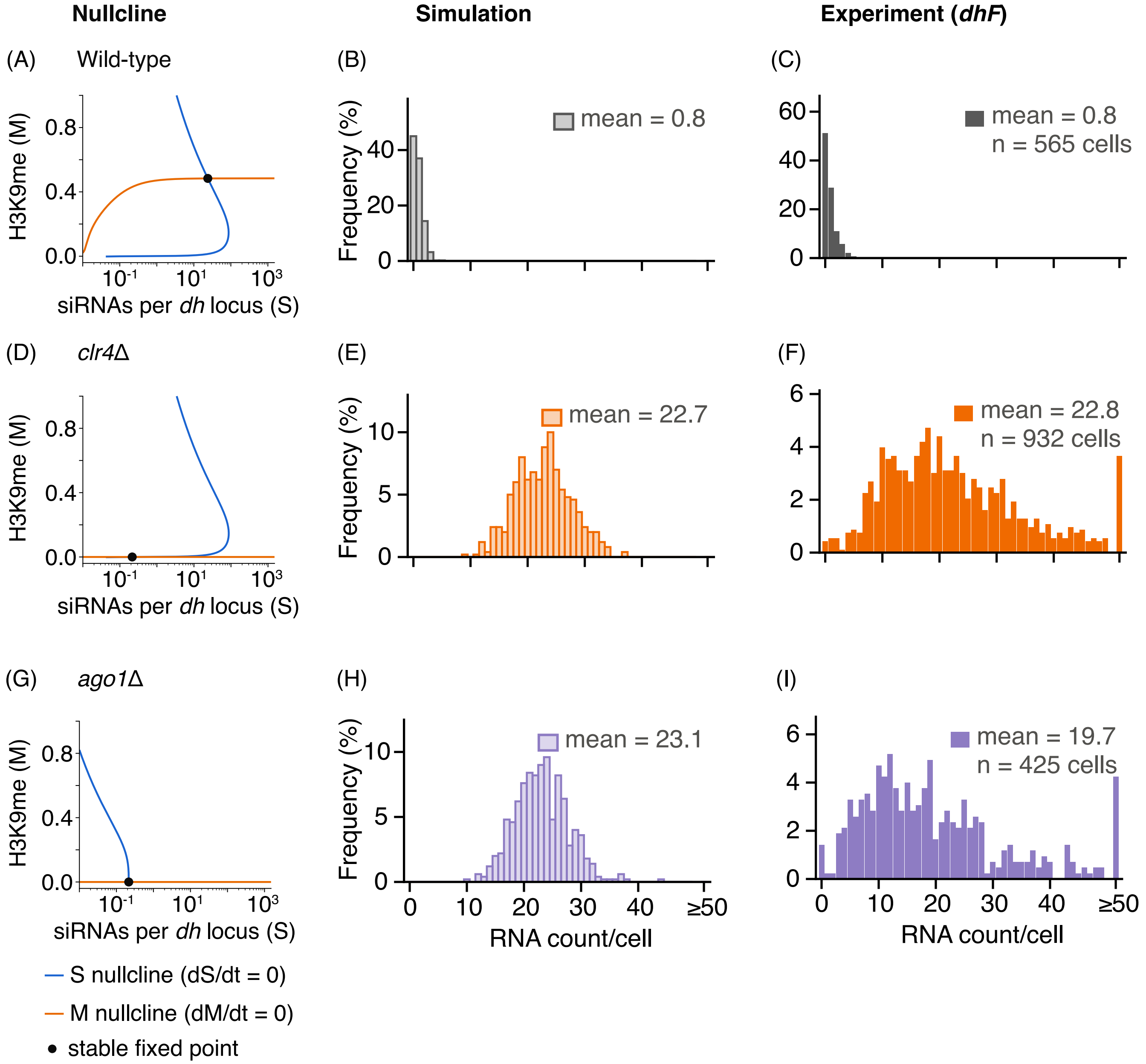

**Supplementary Figure 4: Simulated and experimental transcript abundance histograms for wild-type and mutant cells**

**(A)** Nullcline based on deterministic equations show there is only one steady state in wild-type cells (silenced). Steady-state values for 10 loci together: R* = 0.5, S* = 238, M* = 0.48.

**(B)** RNA histograms obtained from 500 simulation traces for wild-type cells.

**(C)** *dh* forward RNA histograms obtained from smFISH experiments for wild-type cells.

**(D)** Nullcline based on deterministic equations show there is only one steady state in *clr4Δ* cells (desilenced). Steady-state values for 10 loci together: R* = 22.9, S* = 2.2, M* = 0.

**(E)** RNA histograms obtained from 500 simulation traces for *epe1Δ* cells.

**(F)** *dh* forward RNA histograms obtained from smFISH experiments for *epe1Δ* cells.

**(G)** Nullcline based on deterministic equations show there is only one steady state in *ago1Δ* cells (desilenced). Steady-state values for 10 loci together: R* = 22.9, S* = 2.2, M* = 0 (0 of 200).

**(H)** RNA histograms obtained from 500 simulation traces for *ago1Δ* cells.

**(I)** *dh* forward RNA histograms obtained from smFISH experiments for *ago1Δ* cells.

Supplementary Figure 5: Simulated and experimental data for pericentromeric transcription in *epe1* overexpression and *epe1Δ* cells

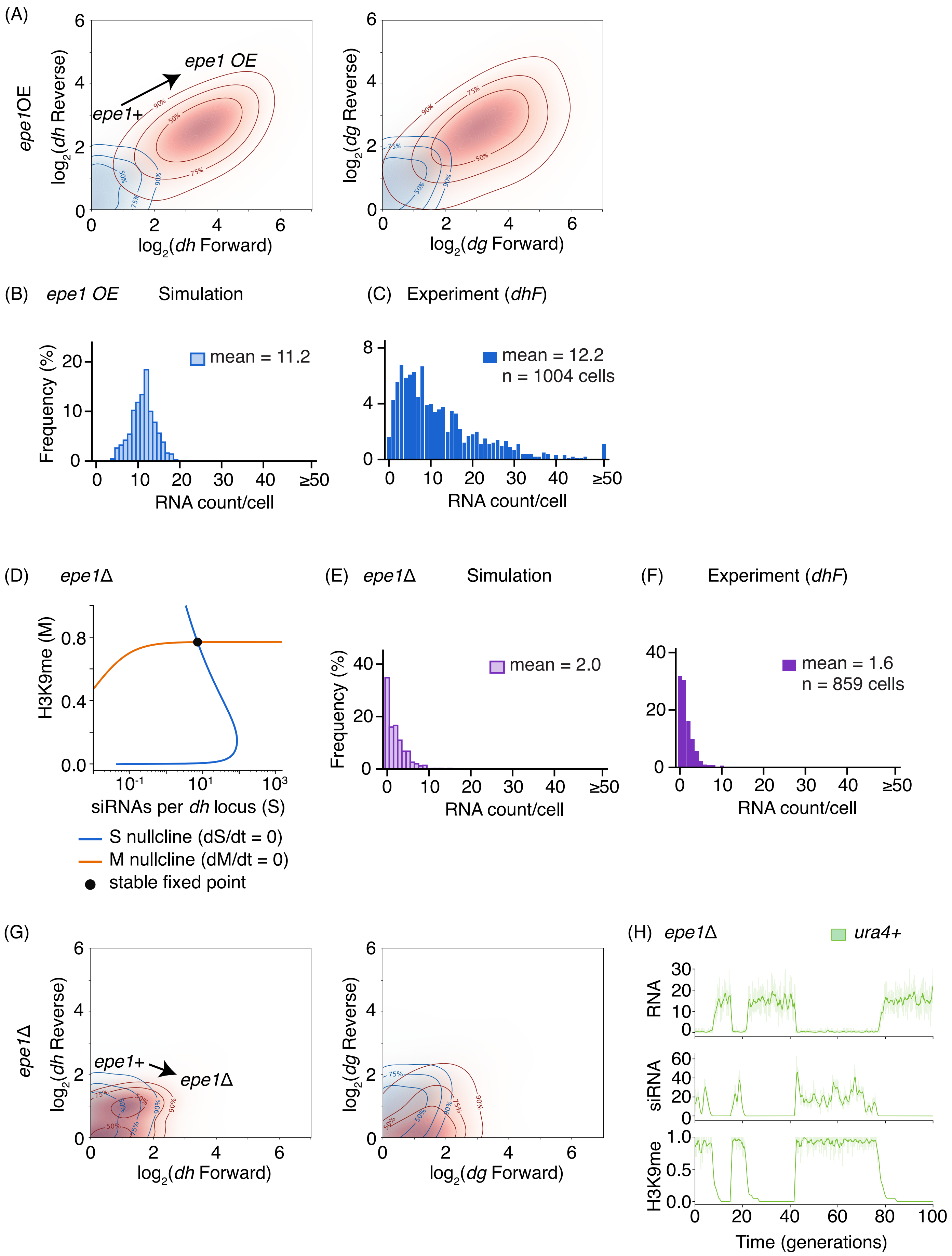

**Supplementary Figure 5: Simulated and experimental data for pericentromeric transcription in *epe1* overexpression and *epe1Δ* cells**

**(A)** Contour plots for *dg* and *dh* transcripts in *epe1* overexpression. Blue and red contours represent the wild-type and the mutant cells respectively.

**(B)** Simulated RNA count per cell histograms obtained from 500 traces for *epe1* overexpression.

**(C)** *dh* forward RNA histograms obtained from smFISH experiments for *epe1* overexpression.

**(D)** Nullcline based on deterministic equations show there is only one steady state at high methylation level in *epe1Δ.* Steady-state values for 10 loci together: $R^* = 0.1$, $S^* = 72$, $M^* = 0.77$.

**(E)** Simulated RNA count per cell histograms obtained from 500 traces for *epe1Δ*.

**(F)** *dh* forward RNA histograms obtained from smFISH experiments for *epe1Δ*.

**(G)** Contour plots for *dg* and *dh* in *epe1Δ*. Blue and red contours represent the wild-type and the mutant cells respectively.

**(H)** Simulation traces for RNA, siRNA, H3K9me for pericentromeric *ura4*+ reporter showing stochastic switching.

Supplementary Figure 6: Flow cytometry controls and replicates for switching of pericentromeric GFP reporter in *epe1Δ* cells

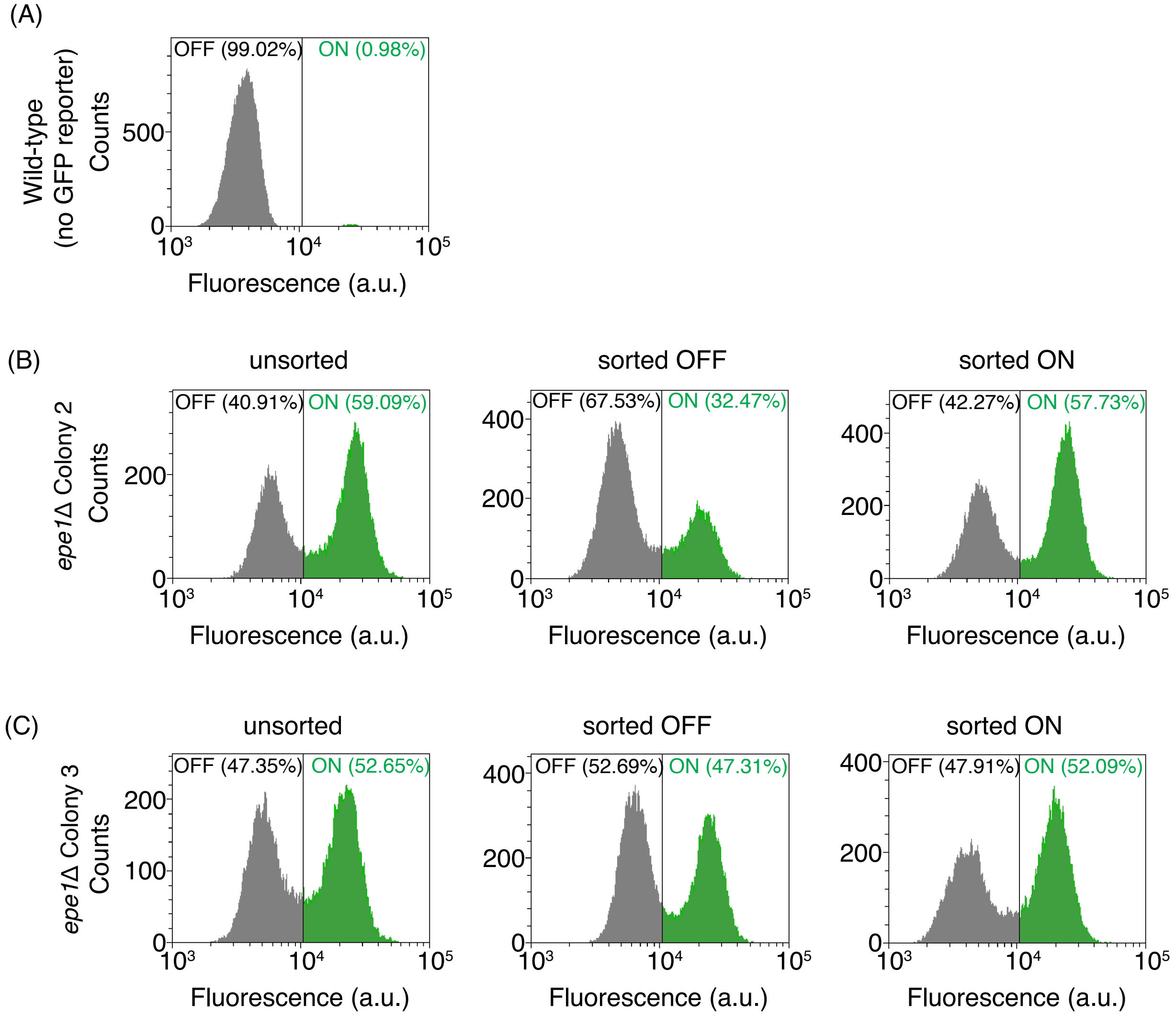

**Supplementary Figure 6: Flow cytometry controls and replicates for switching of pericentromeric GFP reporter in *epe1Δ* cells**

**(A)** Flow cytometry analysis of a strain lacking reporter, used as a negative control.

**(B)** Flow cytometry analysis on *epe1Δ* cells after 4 days of regrowing ON and OFF cells sorted by FACS. Both populations return to a bimodal distribution of ON and OFF cells at steady state.

**(C)** Similar to (B) for cells from another *epe1Δ* colony.
