## Supplementary Note 1 for "Feedback control via H3K9 methylation and siRNA biogenesis autonomously maintains RNAi heterochromatin through continuous low-level transcription"

**SUPPLEMENTARY NOTE 1: Analysis of *dg* and *dh* repeat sequences at the *S. pombe* centromeres**

**Repeat analysis**

In *S. pombe*, *dg* and *dh* are repetitive pericentromeric elements that are present in multiple near-identical copies at the centromeres of all three chromosomes. Here, we discuss the details about RNA-FISH probes used to target these repeat sequences.

All centromeric elements were taken from the PomBase annotations file for the *S. pombe* genome and are named as in PomBase, together with their PomBase IDs (for example, *dg1.1*, *SPRPTCENA.4*; **Table 1**). Throughout the manuscript, the forward transcript is considered to be along the direction of an arrow of that element, and the reverse transcript is in the opposite direction.

We mapped all the annotated repeat sequences along with their orientations (**Supplementary Note 1 Figure 1**). Some stretches within the centromeres are not annotated as *dg* or *dh* repeat elements, though they overlap non-coding RNA genes (**Table 2**). These unannotated stretches were compared by BLAST (Camacho *et al.*, 2009) against all the annotated *dg* and *dh* repeats shown in the figure (**Table 1**). Seven stretches matched an annotated repeat and were labelled *dh*-like or *dg*-like according to the repeat they matched (two on *cen1* and five on *cen3*; none on *cen2*), shaded in **Table 1**. Each one is labelled after its best-matching annotated copy and PomBase ID. In total, there are 20 *dh* and 14 *dg* repeat elements across the three centromeres (34 in total), ranging from 425 to 5,044 bp (*dh*) and from 250 to 6,518 bp (*dg*) in length, respectively. There are 16 *dh* and 11 *dg* annotated in PomBase, plus 4 *dh*-like and 3 *dg*-like stretches identified by BLAST that are not annotated as *dg* or *dh* repeats in PomBase.

**FISH probes and target loci**

The RNA-FISH probes for the *dh* and *dg* regions were designed and purchased from Stellaris Biosciences against the 2,811 bp and 2,001 bp target sequences drawn from *SPRPTCENA.9* and *SPRPTCENA.4*, respectively, on chromosome I. For each target, two sets of 48 probes were designed against the same sequence: one to detect the forward transcript (in the arrow direction; **Tables 2 and 3**) and one to detect the reverse transcript (opposite to the arrow direction; **Tables 4 and 5**).

Because *dg* and *dh* are repetitive pericentromeric elements present in multiple near-identical copies across the centromeres, each probe set was checked computationally for cross-hybridization to other genomic loci beyond its intended target. The mapping was done with the forward-transcript probes; because the reverse-transcript probes were designed against the same targets, the loci identified apply to both sets. The probe sequences were checked against the *S. pombe* genome using the BLAST+ package (Camacho et al., 2009) in a custom-written Python script. A probe was scored as bound at a genomic position only if the alignment had ≤ 1 mismatch (≥ 95% identity for a 20–21 nt oligo). For each probe set, a sliding window equal to the length of its own intended template was scanned along each chromosome. Any window targeted by at least 25 distinct probes was considered a target locus, and all non-overlapping loci were considered final, as any RNA bound by fewer than 25 probes cannot be reliably detected by imaging as per the probe manufacturer. Each probe set targeted 10 loci (**Supplementary Note 1 Figure 1**; **Tables 6 and 7**).

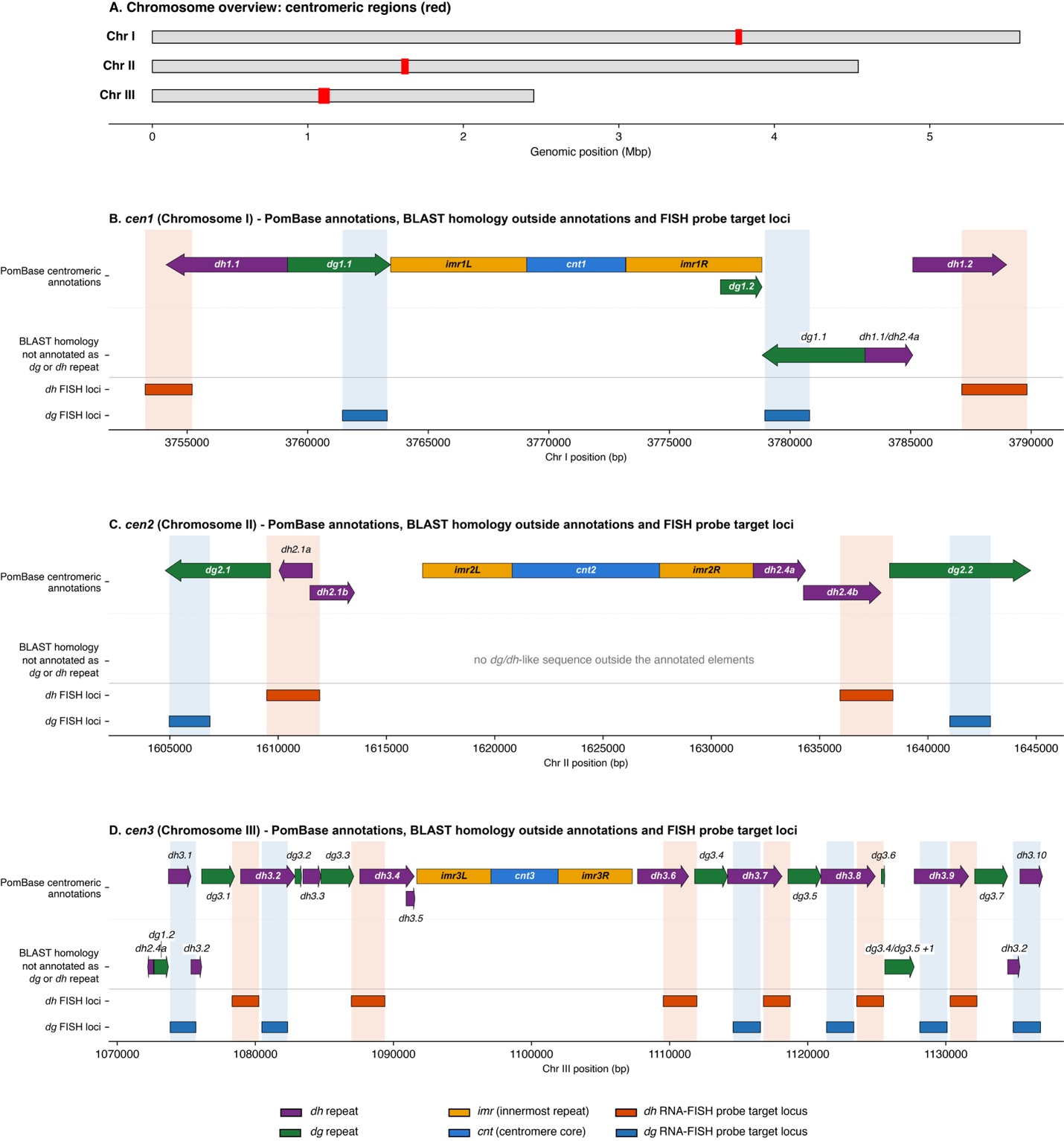

**Supplementary Note 1 Figure 1**: Schematic showing the mapping of *dg* and *dh* sequences across the *S. pombe* genome.

(A) The pericentromeric repeat locations are shown at the genomic scale across all three chromosomes.

(B) *cen1*. Top row: PomBase-annotated *dh* (purple) and *dg* (green) repeats as arrows (the arrow direction is the forward transcript; the opposite direction is the reverse transcript), with *imr* (orange) and *cnt* (blue) as boxes and element names. Middle row: *dg*/*dh*-like stretches found by BLAST that are not annotated as *dg* or *dh* repeats in PomBase, labelled with their best-matching annotated copy (sizes and percentage identity are mentioned in **Table 1**). Bottom rows: loci targeted by *dh* FISH (orange) and *dg* FISH (blue) probes, each in its own row; shaded bands extend each locus through the rows above.

(C) Similar to (B) for centromere 2 (no *dg* or *dh* like sequences found outside the annotated regions).

(D) Similar to (B) for centromere 3.

**Table 1.** ***dh*** **and** ***dg*** **repeats shown in Supplementary Note 1 Figure 1**. Positions are on chromosome I (*cen1*), II (*cen2*) or III (*cen3*); size is estimated as End − Start + 1. Strand is given by PomBase for annotated elements (+ is the arrow direction in the figure, which defines the forward transcript and – denotes the opposite direction). The best-matching identity is obtained from BLAST comparison. Shaded regions show the BLAST-found *dh*-like and *dg*-like stretches that are not annotated as *dg* or *dh* repeats in PomBase.

| **Cen.** | **Element** | **PomBase ID** | **Start (bp)** | **End (bp)** | **Size (bp)** | **Strand** | **Best match (% identity)** |
| --- | --- | --- | --- | --- | --- | --- | --- |
| *cen1* | *dh1.1* | SPRPTCENA.3 | 3,754,127 | 3,759,170 | 5,044 | - |  |
| *cen1* | *dg1.1* | SPRPTCENA.4 | 3,759,165 | 3,763,441 | 4,277 | + |  |
| *cen1* | *dg1.2* | SPRPTCENA.8 | 3,777,115 | 3,778,839 | 1,725 | + |  |
| *cen1* | *dg-like #1* | – | 3,778,840 | 3,783,109 | 4,270 | - | *dg1.1* (99.7%) |
| *cen1* | *dh-like #2* | – | 3,783,110 | 3,785,088 | 1,979 | + | *dh1.1/dh2.4a* (99.9%) |
| *cen1* | *dh1.2* | SPRPTCENA.9 | 3,785,089 | 3,788,981 | 3,893 | + |  |
| *cen2* | *dg2.1* | SPRPTCENB.1 | 1,604,786 | 1,609,649 | 4,864 | - |  |
| *cen2* | *dh2.1a* | SPRPTCENB.3 | 1,610,041 | 1,611,580 | 1,540 | - |  |
| *cen2* | *dh2.1b* | SPRPTCENB.4 | 1,611,467 | 1,613,530 | 2,064 | + |  |
| *cen2* | *dh2.4a* | SPRPTCENB.14 | 1,631,933 | 1,634,352 | 2,420 | + |  |
| *cen2* | *dh2.4b* | SPRPTCENB.15 | 1,634,252 | 1,637,841 | 3,590 | + |  |
| *cen2* | *dg2.2* | SPRPTCENB.17 | 1,638,230 | 1,644,747 | 6,518 | + |  |
| *cen3* | *dh-like #3* | – | 1,072,204 | 1,072,628 | 425 | - | *dh2.4a* (94.5%) |
| *cen3* | *dg-like #4* | – | 1,072,659 | 1,073,707 | 1,049 | + | *dg1.2* (98.4%) |
| *cen3* | *dh3.1* | SPRPTCENC.1 | 1,073,708 | 1,075,339 | 1,632 | + |  |
| *cen3* | *dh-like #5* | – | 1,075,340 | 1,076,114 | 775 | + | *dh3.2* (100.0%) |
| *cen3* | *dg3.1* | SPRPTCENC.2 | 1,076,115 | 1,078,499 | 2,385 | + |  |
| *cen3* | *dh3.2* | SPRPTCENC.4 | 1,078,934 | 1,082,875 | 3,942 | + |  |
| *cen3* | *dg3.2* | SPRPTCENC.5 | 1,082,876 | 1,083,348 | 473 | + |  |
| *cen3* | *dh3.3* | SPRPTCENC.6 | 1,083,449 | 1,084,747 | 1,299 | + |  |
| *cen3* | *dg3.3* | SPRPTCENC.7 | 1,084,748 | 1,087,132 | 2,385 | + |  |
| *cen3* | *dh3.4* | SPRPTCENC.9 | 1,087,567 | 1,091,508 | 3,942 | + |  |
| *cen3* | *dh3.5* | SPRPTCENC.10 | 1,090,919 | 1,091,558 | 640 | + |  |
| *cen3* | *dh3.6* | SPRPTCENC.16 | 1,107,680 | 1,111,378 | 3,699 | + |  |
| *cen3* | *dg3.4* | SPRPTCENC.18 | 1,111,813 | 1,114,198 | 2,386 | + |  |
| *cen3* | *dh3.7* | SPRPTCENC.19 | 1,114,197 | 1,118,138 | 3,942 | + |  |
| *cen3* | *dg3.5* | SPRPTCENC.21 | 1,118,573 | 1,120,957 | 2,385 | + |  |
| *cen3* | *dh3.8* | SPRPTCENC.22 | 1,120,957 | 1,124,898 | 3,942 | + |  |
| *cen3* | *dg3.6* | SPRPTCENC.24 | 1,125,333 | 1,125,582 | 250 | + |  |
| *cen3* | *dg-like #6* | – | 1,125,583 | 1,127,716 | 2,134 | + | *dg3.4/dg3.5/dg3.7* (100.0%) |
| *cen3* | *dh3.9* | SPRPTCENC.25 | 1,127,717 | 1,131,658 | 3,942 | + |  |
| *cen3* | *dg3.7* | SPRPTCENC.27 | 1,132,093 | 1,134,478 | 2,386 | + |  |
| *cen3* | *dh-like #7* | – | 1,134,479 | 1,135,371 | 893 | + | *dh3.2* (96.1%) |
| *cen3* | *dh3.10* | SPRPTCENC.28 | 1,135,372 | 1,137,003 | 1,632 | + |  |

**Table 2. List of *dh* FISH probes targeting the forward transcripts (48 probes).**

| **Probe** | **Sequence (5′→3′)** | **Probe** | **Sequence (5′→3′)** |
| --- | --- | --- | --- |
| 1 | tggaccttgatatggacattg | 25 | ctatgaaatctctacctgctt |
| 2 | atcaagcgagtcgagatgaga | 26 | catgttttggcataaagctgg |
| 3 | gttgcaaaaactaaacgccca | 27 | cgaagttatggagtttcttgc |
| 4 | gatatgagataacccaacgca | 28 | attgatccaacagtatgggta |
| 5 | ttctccaactttgttcaagtt | 29 | tcttgtatctattgctacgtt |
| 6 | aacaacttgcaatatctgcca | 30 | gatatgctcaaaagtgtggcg |
| 7 | tacatttctacaggcacacta | 31 | agtgttgcaagagcgaatact |
| 8 | tccaattgtgatcttccaatg | 32 | caagtggaatgaacatgggca |
| 9 | ccgtatatttaggtttcatgt | 33 | gaacgacaacacatcctgaca |
| 10 | ttgtcctaataggtgcaaagt | 34 | atttttaactcggtggtttct |
| 11 | ctttaatgcagtcacttgagg | 35 | aactgaacaacgcatctacct |
| 12 | tttctgaacctctctgttatg | 36 | tgttggcactttttttctgaa |
| 13 | gtgcggaatgtctacttcaaa | 37 | attggaaggttgtagtgtcag |
| 14 | agcacaagagacatggtgtac | 38 | atgctattccttctttttttt |
| 15 | tgttgtacatttttgcaggac | 39 | tacccatcaactaacgacttg |
| 16 | atgagacaaaggagtccatct | 40 | tagcagctcaactcatgattg |
| 17 | gcattggagatgcgtttaatt | 41 | cgtgacaatacacctaagcta |
| 18 | gtattgtttcagtctcgaagt | 42 | attctatgtcaactgtttgct |
| 19 | tgatgttctttacgaaaccca | 43 | tgattcacgcaaaggatgcat |
| 20 | ttgctttattacacttggtca | 44 | aagtgctctttgtcatgagag |
| 21 | gtttcaaacgactggtagtgt | 45 | caagggaaaaaccgagaccga |
| 22 | aaacaaatcgacttgttgccc | 46 | accatcacagtcttgatttag |
| 23 | gctgcatgtgaattacaacga | 47 | tccgtggacgtatcaactaaa |
| 24 | ccctaaatataacgtgtcgtc | 48 | aagttactcccttgaagtttg |

**Table 3. List of *dg* FISH probes targeting the forward transcripts (48 probes).**

| **Probe** | **Sequence (5′→3′)** | **Probe** | **Sequence (5′→3′)** |
| --- | --- | --- | --- |
| 1 | ggtaccattcaacttggcaa | 25 | attaagcatccttttcgacc |
| 2 | atgacactagtactggtact | 26 | catgaaacgttcaccgtaca |
| 3 | gcctatactgacactagtac | 27 | agatttaagctttcccttga |
| 4 | ccgatactggtgatgatgtt | 28 | gcgggaaaccatttttcttg |
| 5 | tggaagtggtagtggtagtg | 29 | gtgtattcacagaagcatcc |
| 6 | agtggtaagtggaagtggta | 30 | accgctttaacctttcatac |
| 7 | gtggtagtcgtggtagtaat | 31 | gtagttacattcagtgccaa |
| 8 | actccaaagtctttatcctc | 32 | tgcttagcattactttttcc |
| 9 | tagtggcacggccaaaatta | 33 | tgttgcaattgtcccaatta |
| 10 | caggatacctagacgcacaa | 34 | cgctttggcaaaacattgca |
| 11 | ttgcaacgctaaggataccg | 35 | aatttgacgaggcacattcc |
| 12 | tatagtgtgaagccactttc | 36 | gacagtatttgttcagctgg |
| 13 | tgtattagctgtgtcaacca | 37 | gcaaatgcttcacggtattt |
| 14 | actgttcgtgaatgctgaga | 38 | tgagaacttgttatgccaca |
| 15 | acgatatcggttcaatcaca | 39 | ctccatgtacttggaatcga |
| 16 | tgattcttccacatgtttga | 40 | taatccttccttaatcacca |
| 17 | gcagacaatacgtatgcgat | 41 | tgatgttaagggtgaaccgc |
| 18 | acagaagacaagccaagtca | 42 | ttttactacccatgatgtcg |
| 19 | atggattgacattctcgtct | 43 | agtaggaatgatgtactccc |
| 20 | ggaagtgtgattggtcgtac | 44 | tcggttcacttttctctttc |
| 21 | aaagcatttgcactcggttt | 45 | cgtgttgtcgaaacaattcc |
| 22 | ggcacagatcatgattgcta | 46 | tgtttggcatcgccataaat |
| 23 | tcactttgacgagtcttgga | 47 | gtcttggatttatttagcca |
| 24 | gttctctcatcacatcttga | 48 | tattgagatccacagcactc |

**Table 4. List of *dh* FISH probes targeting the reverse transcripts (48 probes).**

| **Probe** | **Sequence (5′→3′)** | **Probe** | **Sequence (5′→3′)** |
| --- | --- | --- | --- |
| 1 | acttcttcacctcatttacaa | 25 | atacaagatatagcgccacac |
| 2 | gcttggttctatgtgttaatt | 26 | ttatcgtcttctttctatacc |
| 3 | tttagccaacattctagttca | 27 | ttatgccaaaacatgcatgca |
| 4 | ttcccttgacaaagctgataa | 28 | taggcatatttcgaagtacca |
| 5 | agttttaactttcttcggtct | 29 | gtgcaaagcaggtagagattt |
| 6 | tctctcatgacaaagagcact | 30 | tgtaattcacatgcagctaca |
| 7 | gcgtgaatcagacacgtagaa | 31 | accagtcgtttgaaacgatga |
| 8 | gaaggtatattgatgcatcct | 32 | atgggtatggacacaagcaac |
| 9 | cgcctatattagcaaacagtt | 33 | ctaatgactttgtgggtttcg |
| 10 | tttagcttaggtgtattgtca | 34 | attaaacgcatctccaatgca |
| 11 | aatcatgagttgagctgctag | 35 | caagtcctgaatcttggcaaa |
| 12 | agggttgttagcctaattatg | 36 | gttgtagttatagttgcagtt |
| 13 | agaaggaatagcataccgtca | 37 | gttgtcttcagagtagtgata |
| 14 | tttcgatggtagtgaagcttt | 38 | cttattgatggcgaagctaga |
| 15 | cactacaaccttccaatcata | 39 | gtttcatcaaagatggactcc |
| 16 | tctttatacgtttgcatgtgt | 40 | aaagtctagtacaccatgtct |
| 17 | gttcagttgactaggatttgt | 41 | tataaatcatcagcctctctc |
| 18 | tttatacatttcccaaggact | 42 | agcataataccgaagcactga |
| 19 | gtcttgtgaaagaaaccaccg | 43 | aacgctatgtttcgtgaatgc |
| 20 | gatgtgttgtcgttcttgaag | 44 | atagtgtgcctgtagaaatgt |
| 21 | ttccacttggatgacagaatc | 45 | acaacatatgcgttgggttat |
| 22 | gaggaaacattcctttgatgc | 46 | gatgaatgggcgtttagtttt |
| 23 | ctcttgcaacactatagttct | 47 | tacaatgattcctctcatctc |
| 24 | tttgttctaagcgaccaagta | 48 | tccatatcaatgtccatatca |

**Table 5. List of *dg* FISH probes targeting the reverse transcripts (48 probes).**

| **Probe** | **Sequence (5′→3′)** | **Probe** | **Sequence (5′→3′)** |
| --- | --- | --- | --- |
| 1 | ggatgttcttccaattgatc | 25 | gtacggtgaacgtttcatgt |
| 2 | actagcattttgcatgcttt | 26 | agagaacaagtcagggtggt |
| 3 | tactcttctcgatgatcctg | 27 | ggttccaagactcgtcaaag |
| 4 | gagtgctgtggatctcaata | 28 | caatcatgatctgtgccttt |
| 5 | tatggcgatgccaaacaaca | 29 | ccgagtgcaaatgcttttgt |
| 6 | ccgattggaattgtttcgac | 30 | atcacacttcctatagctga |
| 7 | ggagtacatcattcctactt | 31 | gtcaatccatcatgtacgac |
| 8 | aagaactttttccatccgca | 32 | gtcttctgtatgtgatccaa |
| 9 | gtttttttccacaacaccag | 33 | tacgtattgtctgctctgac |
| 10 | accaacgacatcatgggtag | 34 | agtttcctgtacatcttcta |
| 11 | cacccttaacatcatgtttt | 35 | gcattcgtttcgcttatgaa |
| 12 | aaacgaccaatatgctgcgg | 36 | tctcagcattcacgaacagt |
| 13 | ggaaggattactgtgcatct | 37 | agtggaaagtggcttcacac |
| 14 | gtacatggagagcgtatgtt | 38 | gtatttgtgcgtctaggtat |
| 15 | tgtggcataacaagttctca | 39 | aaataattttggccgtgcca |
| 16 | ccgtgaagcatttgctttat | 40 | ctacgatttcttgttagcga |
| 17 | cccagctgaacaaatactgt | 41 | gcttgttgtacgttgttcaa |
| 18 | tgtgcctcgtcaaattatca | 42 | catctccattcttgtcatta |
| 19 | attgtatctttcgtttgcgt | 43 | cgactaccactaaccatcac |
| 20 | tgcaatgttttgccaaagcg | 44 | cttaccacttccattactac |
| 21 | ttaaagcggttgtttggcac | 45 | aacatcatcaccagtatcgg |
| 22 | gtggatgcttctgtgaatac | 46 | gtactagtgtcagtataggc |
| 23 | aagaaaaatggtttcccgcc | 47 | accagtactagtgtcatcta |
| 24 | ttcattgaatcacctttcca | 48 | ccaagttgaatggtaccatt |

**Table 6. List of loci targeted by *dh* FISH probes (≥ 25 of 48 probes, ≤ 1 mismatch).**

| **S. No.** | **Cen.** | **Start (bp)** | **End (bp)** | **Width (bp)** | **Probes bound (/48)** |
| --- | --- | --- | --- | --- | --- |
| 1 | *cen1* | 3,753,263 | 3,755,213 | 1,951 | 37 |
| 2 | *cen1* | 3,787,122 | 3,789,825 | 2,704 | 48 |
| 3 | *cen2* | 1,609,474 | 1,611,920 | 2,447 | 45 |
| 4 | *cen2* | 1,635,942 | 1,638,388 | 2,447 | 45 |
| 5 | *cen3* | 1,078,318 | 1,080,256 | 1,939 | 33 |
| 6 | *cen3* | 1,086,951 | 1,089,382 | 2,432 | 44 |
| 7 | *cen3* | 1,109,543 | 1,111,976 | 2,434 | 44 |
| 8 | *cen3* | 1,116,796 | 1,118,736 | 1,941 | 33 |
| 9 | *cen3* | 1,123,556 | 1,125,496 | 1,941 | 33 |
| 10 | *cen3* | 1,130,316 | 1,132,256 | 1,941 | 33 |

**Table 7. List of loci targeted by *dg* FISH probes (≥ 25 of 48 probes, ≤ 1 mismatch).**

| **S. No.** | **Cen.** | **Start (bp)** | **End (bp)** | **Width (bp)** | **Probes bound (/48)** |
| --- | --- | --- | --- | --- | --- |
| 1 | *cen1* | 3,761,442 | 3,763,298 | 1,857 | 48 |
| 2 | *cen1* | 3,778,957 | 3,780,812 | 1,856 | 47 |
| 3 | *cen2* | 1,604,972 | 1,606,852 | 1,881 | 47 |
| 4 | *cen2* | 1,641,011 | 1,642,891 | 1,881 | 47 |
| 5 | *cen3* | 1,073,831 | 1,075,696 | 1,866 | 48 |
| 6 | *cen3* | 1,080,472 | 1,082,337 | 1,866 | 48 |
| 7 | *cen3* | 1,114,599 | 1,116,581 | 1,983 | 48 |
| 8 | *cen3* | 1,121,359 | 1,123,341 | 1,983 | 48 |
| 9 | *cen3* | 1,128,119 | 1,130,101 | 1,983 | 48 |
| 10 | *cen3* | 1,134,879 | 1,136,861 | 1,983 | 48 |
