## Supplementary Note 2 for "Feedback control via H3K9 methylation and siRNA biogenesis autonomously maintains RNAi heterochromatin through continuous low-level transcription"

**SUPPLEMENTARY NOTE 1: Modeling RNAi-mediated gene silencing**

RNAi-mediated heterochromatin formation in fission yeast involves three key components: RNA Pol II transcription of the target locus, processing of the nascent transcripts into small interfering RNAs (siRNAs) by the RDRC/Dicer1 machinery, and RITS-mediated establishment of H3K9 methylation, which in turn represses transcription (**Figure 4A**). Here, we develop a simple mathematical framework building on earlier theoretical work describing heterochromatin systems (Dodd *et al.*, 2007; Nickels *et al.*, 2021; Abdulla, Vaillant and Jost, 2022; Ghimire, Motamedi and Joh, 2024) and informed by our perturbation experiments to characterize the self-reinforcing loop between RNA interference (RNAi) and H3K9 methylation that maintains heterochromatin in fission yeast.

**Model overview**

We consider a genomic locus with N = 20 nucleosomes (~3kb), analogous to a single *dh* repeat of the pericentromeric region in *S.pombe* (non-coding; the *dh* repeat transcripts are produced from 10 locations) or to the *ura4* gene placed in the pericentromere as a reporter (coding) and follow the dynamics of Pol II-derived RNA R, small interfering RNA (siRNA) S, and the chromatin state (M) of the locus. The chromatin state is defined as the fraction of nucleosomes methylated, M = m/N where m represents the number of methylated nucleosomes.

The following system of equations describes the dynamics:

$\frac{\mathrm{dR}}{\mathrm{dt}}=\alpha H_{1}(M)-\gamma RMH_{2}(S)-\delta_{1}R$ (S1)

$\frac{\mathrm{dS}}{\mathrm{dt}}=n_{S}\gamma RMH_{2}(S)+\nu R-\delta_{2}S$ (S2)

$\frac{\mathrm{dM}}{\mathrm{dt}}=RH_{2}(S)(\varepsilon_{\mathrm{nuc}}+\varepsilon_{s}M)(1-M)-\delta_{3}M$ (S3).

Transcripts are produced at a rate α when the locus is not methylated where α is the output of one copy of the locus, in transcripts per minute (the 10 *dh* copies are simulated as independent copies and their transcripts summed per cell; *ura4* is a single copy). The Hill function $H_{1}\left( M \right)=\frac{1}{1+\left( \frac{M}{K} \right)^{\rho}}$ represents methylation based transcriptional silencing of the locus, wherein the genomic region of interest is silent when M > K. K is the methylation fraction that halves transcription, and the Hill coefficient $\rho$ (=3, a specific choice that doesn’t affect the general conclusions) sets how switch-like repression is.

siRNAs are produced in two ways. The main source is cleavage by RITS complex which requires both siRNA guides and H3K9 methylation to hold RITS on chromatin (Chp1). This process describes the co-transcriptional silencing mechanism. We write this cleavage as a term proportional to the number of transcripts and the methylation fraction, and saturating in the number of siRNAs ($n_{S}\gamma RMH_{2}(S)$), where $H_{2}(S)=S/(K_{S}+S)$ and $K_{S}$ is the siRNA level giving half-maximal activity. This source amplifies the siRNAs already present. *De novo* or basal (primal) siRNA production from transcripts happens at a much slower rate ($\gamma\gg\nu$) independent of methylation state (Halic and Moazed, 2010). Each cleavage event converts one transcript into $n_{S}$ = 10 siRNAs.

Deposition of methylation by histone methyltransferase, Clr4 has two components:
(i) nucleation, siRNA-guided RITS bound to a transcript recruits Clr4 through the adaptor Stc1 (Bayne *et al.*, 2010). This step requires transcripts and siRNAs but not pre-existing methylation, thus it can *de novo* establish a heterochromatin domain.
(ii) spreading, that requires existing methylation. RITS binds methylated nucleosomes through the chromodomain of its Chp1 subunit(Motamedi *et al.*, 2004). Existing methylation enhances the activity of Clr4 $(\varepsilon_{s}\gg\varepsilon_{\mathrm{nuc}})$ (Al-Sady, Madhani and Narlikar, 2013; Newar *et al.*, 2022) For the sake of simplicity, spreading is gated by the same transcript and siRNA dependent factor as nucleation, $RH_{2}(S)$, and is proportional to M.

The first order decay of RNA ($\delta_{1}$) is set to give an RNA half-life of 10-20 mins(Eser *et al.*, 2016), and the cleavage mechanism ($\gamma$) acts on a similar or faster timescale at a methylated locus. siRNA are lost at a first-order rate ($\delta_{2}$) that combines degradation and dilution by growth, most siRNAs are degraded within a few hours after their production is shut off(Pisacane and Halic, 2017). Methylation is lost at a first-order rate ($\delta_{3}$). This combines active removal promoted by the protein Epe1, histone turnover (Ragunathan, Jih and Moazed, 2015), and dilution of the marks when new histones are deposited at replication. Since both nucleation and spreading require transcripts and siRNAs, methylation at this locus always depends on RNAi based deposition in this model. Removing Epe1 lowers the loss rate; Clr4-tethering at ectopic locus identifies Epe1 as the main barrier to RNAi-independent inheritance of H3K9 (Ragunathan, Jih and Moazed, 2015).

The formalism describes gene silencing as a combined effect of transcriptional repression with cleavage of RNA that is still made. A property of this structure is that siRNAs amplify themselves, a steady state with siRNAs present requires amplification to nearly balance the loss: $n_{S}\gamma RM/(K_{S}+S)\approx\delta_{2}$ (the basal term νR being small). A silent locus therefore holds the product of its transcript level and its methylation level close to $\delta_{2}(K_{S}+S)/(n_{S}\gamma)$, which is set by the siRNA level. When methylation input is weakened, siRNA levels rise and partly compensate. This is why mutants with weakened methylation (*clr4 W31G*) can stay largely silent.

**Simulations**

For a given set of parameters, the stochastic dynamics of the system is simulated using the Gillespie algorithm implemented on Python using GillesPy2 (Matthew *et al.*, 2023). R, S and the number of methylated nucleosomes m are integers, the terms of the equations are the reaction propensities, and each cleavage event converts one R into $n_{S}$ = 10 siRNAs. Starting from a silent state (R = 0, S = 20, M = 0.9), the system relaxes to a steady state. Each simulation corresponds to a ‘single cell’ trajectory of the cell state defined by R, M and S; for the non-coding set R, M and S are summed across 10 trajectories, one for each *dh* copy. In each condition, 500 different trajectories of 40 generations (generation time 150 min) were simulated and the steady-state values are shown; time traces show 100 generations after waiting for 20 generations to let the trajectories reach the steady state. Mutants are simulated by changing the wild-type parameters: *clr4Δ*, $\varepsilon_{\mathrm{nuc}}=\varepsilon_{s}=0$; *ago1Δ*, $\gamma=\varepsilon_{\mathrm{nuc}}=\varepsilon_{s}=0$; *clr4*-W31G, $\varepsilon_{s}=0$ (nucleation only); *epe1Δ*, no Epe1-dependent removal of methylation ($\delta_{3}=\mu$) where $\mu$ represents the cell-cycle dependent dilution of H3k9me; *epe1*-OE, Epe1-dependent removal increased from 0.05 to 2.5 min⁻¹ ($\delta_{3}=2.5+\mu$).

**Steady state and nullclines**

For analytic steady state, we set the transcript (fastest dynamics) equation to zero which gives R as a function of M and S:

$$R^{*}(M,S)=\frac{\alpha H_{1}(M)}{\delta_{1}+\gamma MH_{2}(S)}$$

Substituting R* reduces the steady-state problem to two curves in the plane of methylation and siRNA, the nullclines. Along the siRNA nullcline S solves a quadratic with one positive root:

$$\delta_{2}(\delta_{1}+\gamma M)S^{2}+[\delta_{2}\delta_{1}K_{S}-\alpha H_{1}(M)(n_{S}\gamma M+\nu)]S-\alpha H_{1}(M)\nu K_{S}=0$$

Along the methylation nullcline:

$$S=\frac{K_{S}\delta_{1}\delta_{3}M}{\alpha H_{1}(M)A-\delta_{3}M(\delta_{1}+\gamma M)}, A=(1-M)(\varepsilon_{\mathrm{nuc}}+\varepsilon_{s}M)$$

Every crossing of the two curves is a steady state of the full model. This reduction is exact at steady state; it was used only to locate steady states, and all the time courses integrate the full three-variable model. Stability was assessed from the eigenvalues of the full 3 × 3 Jacobian, evaluated by finite differences. A steady state is stable when all eigenvalues have negative real parts. The nullclines of wild-type, *clr4Δ* and *ago1Δ* show single stable fixed point as expected, a silenced state with high methylation in wild-type and a desilenced state without methylation in *clr4Δ* and *ago1Δ* (**Supplementary Figures 4A,D,G**).

**Parameter values**

We constrain most of the parameters in the equations based on the experimental data for *dh* Forward strand (*dhF*) and previously reported timescales, with five free parameters in the model $\gamma, \nu, K_{S}, \varepsilon_{\mathrm{nuc}}$ and $\varepsilon_{s}$. Using the observed *clr4Δ* RNA-FISH mean of ~20 RNA/cell together with δ_1_ (from the measured RNA half-life) gives α ~ 1.5 RNA/min for the ten *dh* copies (0.15 RNA/min per copy).

| **Symbol** | **Non-coding** | **Units** | **Basis** |
| --- | --- | --- | --- |
| α | 0.15 | RNA min⁻¹ | Transcription rate of one unmethylated copy of the locus. Constrained jointly with δ_1_ by the *clr4Δ* smFISH mean (~20 RNA/cell). |
| δ_1_ | 0.0656 | min⁻¹ | Basal RNA decay (0.06 min⁻¹(Eser *et al.*, 2016) plus dilution by cell division and conversion to primal siRNA. |
| δ_2_ | 0.0106 | min⁻¹ | siRNA degradation rate 0.006 min⁻¹(Pisacane and Halic, 2017) plus dilution by cell division. |
| δ_3_ | 0.0546 | min⁻¹ | Rate of methylation turnover: Epe1-dependent removal at 0.05 min⁻¹ (0 in *epe1Δ*, 2.5 in *epe1*-OE) plus dilution by cell division. |
| γ | 1 | min⁻¹ | siRNA biogenesis rate |
| ν | 10⁻³ | min⁻¹ | Primal siRNA production rate |
| K | 0.3 | – | Methylation fraction that halves transcription |
| ρ | 3 | – | Hill coefficient of transcriptional repression |
| K_S_ | 1 | siRNAs | siRNA level for half-saturated cleavage and methylation |
| n_S_ | 10 | siRNAs | siRNAs produced per cleavage event |
| ε_nuc_ | 0.01 | min⁻¹ | Nucleation by RITS recruitment of Clr4 |
| ε_s_ | 2 | min⁻¹ | Spreading from existing methylation |

For *ura4+*, α = 1 RNA/min gives ~15 RNA/cell in *clr4Δ* (check against *ura4+* RNA-FISH). Compared with the non-coding RNA (*dhF*), the parameter set for coding RNA (*ura4*) has these changes: γ = 0.2, K_S_ = 30 and ν= 2×10⁻⁵, and one copy of the locus is simulated instead of ten.

**Results**

The Stochastic simulations (Gillespie) of wild-type cells converge onto a stable fixed point where non-coding RNA transcripts are present at low levels (~1 per cell), while siRNA and H3K9me are constant at high levels (**Figure 4C**). To systematically characterize the model, we analyze how the free parameters affect the system. The phase diagrams of γ vs ε_nuc_ and γ vs ε_s_ show a range of parameter values over which the model reproduces the expected RNA levels of the genotypes under consideration (**Figure 4E**).

The mean RNA values obtained from simulations agree with the experimental RNA-FISH measurements for all the genotypes under consideration (**Supplementary Figures 4C,F,I & 5C,F**). In wild-type, the siRNA level is about S = 450/cell and M ~ 0.46. In *clr4Δ* and *ago1Δ*, there is no siRNA amplification, so R value is α/δ_1_ (~ 23 RNA per cell for *dh*). In *clr4 W31G*, methylation is low (M ~ 0.12) and the RNA levels rise compared to wild-type but maintained at lower levels compared to *clr4Δ*.

To test how demethylation affects the silenced state, we varied the Epe1-dependent removal srate alone for *dh* (0 in *epe1Δ*, 0.05 min^-1^ in WT, 2.5 min^-1^ in *epe1*-OE). The methylation fraction decreases as the rate increases and it is highest in *epe1Δ* (**Figure 5A**). The percentage of cells with RNA >3 is low in wild-type and rises when the demethylation rate is either reduced or increased (**Figure 5B**). The siRNA level increases as the removal rate rises from zero, continues to rise through the wild-type value, peaks at a higher rate before falling off to low levels (**Figure 5C**). The *epe1Δ* RNA traces show a huge variability in RNA levels over time compared with other genotypes, showing that Epe1-based methylation turnover is needed for stable RNA levels, otherwise the locus switches transiently to a desilenced state. The nullcline for *epe1Δ* shows a single steady state (**Supplementary Figure 5D**), so the switching to high RNA levels seen is not due to bistability. siRNA levels are low in *epe1Δ* compared to wild-type. Switching is clearly demonstrated in experiments with our single-locus reporter gene *ura4*, so we simulated its traces for *epe1Δ* (**Supplementary Figure 5H**), using the *ura4+* (coding) gene parameters for transcription (α =1) and RNAi pathway (γ = 0.2, K_S_ = 30 and ν= 2×10⁻⁵). These traces show that cells maintain low levels of siRNA that are driven to extinction by stochastic fluctuations. Without siRNA, there is no reinforcement of heterochromatin, and methylation marks are gradually diluted by cell division reaching an actively transcribing state but the zero siRNA, zero methylation state is not absorbing because of the basal siRNA production ν which sets the timescale of transcribing to silent switch. Therefore, stable silencing requires siRNA levels to be maintained, which in turn requires a low level of transcription.

Overall, these simulations show that a feedback loop between RNA, siRNA and methylation status of the chromatin domain, balanced by Epe1-dependent removal of methylation marks, explains the RNAi-regulated silencing of *S.pombe* pericentromeric repeats. The parameters generate a stable steady state which is not completely silent but sets transcriptional set point and this may be a general feature of RNA-based silencing architectures.
