## Supplementary Table 1 for "Feedback control via H3K9 methylation and siRNA biogenesis autonomously maintains RNAi heterochromatin through continuous low-level transcription"

**Supplementary Table 1. *S. pombe* strains used in this study.**

| **KR Strain #** | **Genotype** | **Source** | **Figure(s)** |
| --- | --- | --- | --- |
| 327 | *h-* *leu1-32* | Moazed lab | Figure 1C; Supplementary Figure 6A |
| 3972 | h- *ura4::padh1+-Fragment 1(barcode)-ter-Bboxes-natMX6* *cnp1-mNeonGreen-sfGFP-hphMX6* | This study | Figure 1D, 2I |
| 1 | *h+* *leu1-32* *ade6-216* *ura4-D18* *imr1R(Ncol)::ura4+* *oriI* *ura4Δ::nmt1-APEX2-ADht* | Moazed lab | Figure 1F |
| 3981 | *h+* *otr1R(SphI)::ura4+* *ura4-D18* *leu1-32* *ade6-M210* *cnp1-mNeonGreen-sfGFP-hphMX6* | This study | Figure 1G, 2A-C, 3J, 5F; Supplementary Figure 2A-D, 4C |
| 963 | *h+* *leu1* *ura4* *ade6-210* *IRC1R::ura4* | Jia lab | Figure 1H |
| 4034 | *h+* *leu1-32* *ade6-210* *ura4Δ::bsdMX6* *otr1R(SphI)::ura4+* *clr4Δ:KanMx6* *cnp1-mNeonGreen-sfGFP-hphMX6* | This study | Figure 1I, 2A-C, 3J, 5F; Supplementary Figure 2A, 2B, 2E, 4F |
| 4524 | h+ *leu1-32* *ade6-M210* *otr1R(SphI)::ura4+* *ura4Δ::bsdMX6* a*go1Δ::kanMX6* | This study | Supplementary Figure 1F |
| 3528 | *h+ leu1-32* *ade6-M210* *otr1R(SphI)::ura4+* *ura4-DS/E ago1Δ::kanMX6* | Moazed lab | Figure 2D, 3J; Supplementary Figure 2F, 4I |
| 3531 | *h+ leu1-32* *ade6-M210* *otr1R(SphI)::ura4+* *ura4-DS/E swi6Δ::natMX6* | Moazed lab | Figure 2E; Supplementary Figure 2I |
| 3976 | *h-* *ura4::padh1+-Fragment 1(barcode)-ter-Bboxes-natMX6* *clr4Δ::kanMX6* *cnp1-mNeonGreen-sfGFP-hph* | This study | Figure 2I |
| 3529 | *h+ leu1-32* *ade6-M210* *otr1R(SphI)::ura4+* *ura4-DS/E dcr1Δ::natMX6* | Moazed lab | Supplementary Figure 2G |
| 3303 | *h+* *leu1-32* *ade6-M210* *otr1R(SphI)::ura4+* *ura4Δ::bsdMX6* *rdp1Δ::TAP-kanMX6* | This study | Supplementary Figure 2H |
| 3331 | *h+* *leu1-32* *ade6-M210 otr1R(SphI)::ura4+* *ura4-D18* *ppcn1-eGFP-pcn1-bsdMX6* | This study | Figure 3B-G; Supplementary Figure 3A-C |
| 3418 | *h+* *leu1-32* *ade6-M210* *otr1R(SphI)::ura4+* *ura4-D18* *cdc2-asM17-bsdMX6* *ppcn1-eGFP-pcn1-bsdMX6* | This study | Figure 3G, 3H; Supplementary Figure 3D-F |
| 4257 | *h+* *leu1-32* *ade6-M210 otr1R(SphI)::ura4+* *ura4-D18*, *ppcn1-eGFP-pcn1-bsdMX6*, *clr4 R406H* | This study | Supplementary Figure 3G-N |
| 3377 | *h+* *leu1-32* *ade6-M210 otr1R(SphI)::ura4+* *ura4-D18*, *ppcn1-eGFP-pcn1-bsdMX6* *clr4 W31G* | This study | Figure 4F-H |
| 1267 | *h-* *ade6::ade6+-adh15prom-skp1-OsTIR1-natMX6-adh15prom-skp1-AtTIR1-2NLS* *ura4-D18* *nmt41-epe1-3xFLAG-AID-kanMX6* | Larkin, et.al, Dev Cell. 2024 | Figure 5E, 5F; Supplementary Figure 5A,C |
| 3309 | *h+* *leu1-32* *ade6-M210* *otr1R(SphI)::ura4+* *ura4∆::bsdMX6* *epe1Δ::kanMX6* | This study | Figure 5H, 5I; Supplementary Figure 5F, 5G |
| 4485 | SPY137 *h+* *otr1R(SphI)::pcam1-mNeonGreen-GFP* *ura4∆::bsdMX6* *leu1-32* *ade6-M210* *epe1Δ::kanMX6* | This study | Figure 6A-C,E; Supplementary Figure 6B, 6C |
| 4509 | *h+* *leu1-32* *ade6-210* *ura4Δ::bsdMX6* *otr1R(SphI)::pcam1-mNeonGreen-GFP* *clr4Δ::kanMX6* | This study | Figure 6A |
| 4546 | *h-* *leu1-32* *ade6-210*? *ura4Δ::bsdMX6 otr1R(SphI)::pcam1-mNeonGreen-GFP* | This study | Figure 6A |
